# Maternal SMARCA5 is required for major ZGA in mouse embryos

**DOI:** 10.1101/2023.12.05.570276

**Authors:** Oana Nicoleta Kubinyecz, Deborah Drage, Fatima Santos, Christel Krueger, Hanneke Okkenhaug, Celia Alda Catalinas, Melanie Eckersley-Maslin, Jasmin Taubenschmid-Stowers, Wolf Reik

## Abstract

Zygotic genome activation (ZGA) in mice takes place in two waves, a minor wave in the one-cell embryo, and a major wave at the two-cell stage, both accompanied by global transcriptional and epigenetic reprogramming. However, the orchestration of these reprogramming events by maternal factors deposited in the oocyte is not yet entirely understood. We and others have recently shown that epigenetic modifiers such as SMARCA5 (the main ATPase in ISWI complexes) can initiate the ZGA transcriptional programme *in vitro*. So far, the role of SMARCA5 in ZGA *in vivo* has not been addressed, as constitutive knock-out mice lacking SMARCA5 are not viable. We have overcome this limitation by using the targeted protein-depletion system Trim Away to degrade SMARCA5 in early zygotes. We further harnessed the power of single cell multi-omics (scNMT-seq) and showed that in the absence of SMARCA5, major ZGA genes fail to be upregulated at the two-cell stage. This is explained by the lower accessibility and disrupted nucleosome positioning at their promoters and distal regulatory regions, compared to wild-type embryos. In contrast, we show that global chromatin accessibility at the two-cell stage is higher in SMARCA5 depleted embryos compared to control embryos, and this is accompanied by other global structural changes involving heterochromatic regions. Our results show that SMARCA5 has a global regulatory role at the two-cell stage, which includes the control of ZGA gene promoters and distal regulatory regions.

## Introduction

The journey of all new mammalian organisms starts with the moment of fertilisation and the formation of a one-cell embryo (zygote), containing haploid maternal and paternal nuclei (pronuclei). At this stage, the two pronuclei need to undergo epigenetic reprogramming in order to be able to give rise to a totipotent zygote, as part of the complex maternal to zygotic transition (MZT) (Eckersley-Maslin et al., 2018). In mouse embryos, the first transcriptional events take place in the zygote, followed by a major transcriptional burst at the two-cell stage (K. Abe et al., 2018; K. -i. Abe et al., 2015; Yamamoto et al., 2016). This genome awakening event is known as zygotic genome activation (ZGA), comprising the minor and the major ZGA in the zygote and the two-cell stage, respectively (reviewed in Eckersley-Maslin et al., 2018).

To study early developmental processes such as ZGA *in vitro*, stem cell model systems have been used. So called mouse 2C-like cells (2CLCs) (Macfarlan et al., 2012) and human 8C-like cells (Taubenschmid-Stowers et al., 2022) exist among embryonic stem cells (ESCs) that transcriptionally and epigenetically resemble ZGA stage embryos. 2CLCs have been widely used to identify candidate regulators of genome activation in a high-throughput way *in vitro* and one such screen identified the chromatin remodeler SMARCA5 as an inducer of ZGA-like transcription in mouse ESCs (Alda-Catalinas et al., 2020; Rodriguez-Terrones et al., 2018; Zhao et al., 2018). SMARCA5 (SWI/SNF-related matrix-associated actin-dependent regulator of chromatin subfamily A member 5, or Snf2h) is the main ATPase in ISWI complexes that slide histones along the DNA double strand and act as a ruler for nucleosome positioning and distribution. SMARCA5 deficient ESCs are viable, however CTCF binding was disturbed in those cells and differentiation impaired (Barisic et al., 2019; Wiechens et al., 2016).

SMARCA5 plays a crucial role during oocyte development, and it is deposited in the mature oocytes both as protein and mRNA (Zhang et al., 2020). It has been shown to be lethal during peri-implantation in mice in various studies: *Smarca5* knockout embryos fail to develop normal inner cell mass (ICM) structures in blastocysts (Stopka & Skoultchi, 2003) and *Smarca5* siRNA-mediated knockdown embryos fail to develop past the 8-cell (Torres-Padilla & Zernicka-Goetz, 2006) or morula to blastocyst stage (Shi et al., 2021). Moreover, absence of SMARCA5 in oocytes leads to metaphase II (MII) stage defects (Zhang et al., 2020), and this made it difficult to study the role of (maternally deposited) SMARCA5 in zygotes and early embryos.

We asked if maternal SMARCA5 is required for mouse ZGA *in vivo*, and for this we used maternal knockout mouse models and the Trim-Away protein degradation system (Clift et al., 2017) in combination with single-cell nucleosome, methylation, and transcription sequencing (scNMT-seq) (Clark et al., 2018) to assess the gene expression and epigenetic changes in SMARCA5 depleted embryos. We found that SMARCA5 deficient embryos fail to express two-cell specific transcripts and endogenous retroviral elements and thus established a requirement for SMARCA5 in mouse ZGA *in vivo*. We further discovered that chromatin at putative regulatory regions (which we defined as open chromatin regions in the two-cell stage embryos that are not promoters) and at ZGA gene promoters in two-cell embryos (but not other promoters) fail to open locally in absence of SMARCA5. These changes are opposite to the increased global chromatin accessibility induced by the absence of SMARCA5. Moreover, heterochromatin regions around nucleolus precursor bodies that require SMARCA5 fail to organise correctly in the two-cell stage embryos, as observed by the mislocalization of HP1-β.

## Results

### Maternally deposited SMARCA5 is crucial for preimplantation development beyond the two-cell stage

We first assessed the presence of SMARCA5 in wild-type (CB57L/6) embryos by immunofluorescence and confirmed its nuclear localisation in oocytes and throughout preimplantation development (Fig. 1A) (Shi et al., 2021). In order to test if maternally deposited SMARCA5 is required for mouse ZGA, we used *Smarca5* conditional knockout mice (*Smarca5^fl/fl^*) in which exon 5 of the gene is flanked by *loxP* sites (Alvarez-Saavedra et al., 2019). These mice were crossed with *Zp3-Cre* positive males to generate *Smarca5^fl/fl^; Zp3-Cre* females (Fig. S1A) that give rise to maternal knockout oocytes and embryos (*Smarca5 matKO*), lacking the maternal SMARCA5 deposits, while having a wild-type paternal allele. We confirmed the absence of SMARCA5 in germinal vesicle (GV) oocytes derived from *Smarca5^fl/fl^; Zp3-Cre* females by immunofluorescence (Fig. S1B).

**Figure 1.**
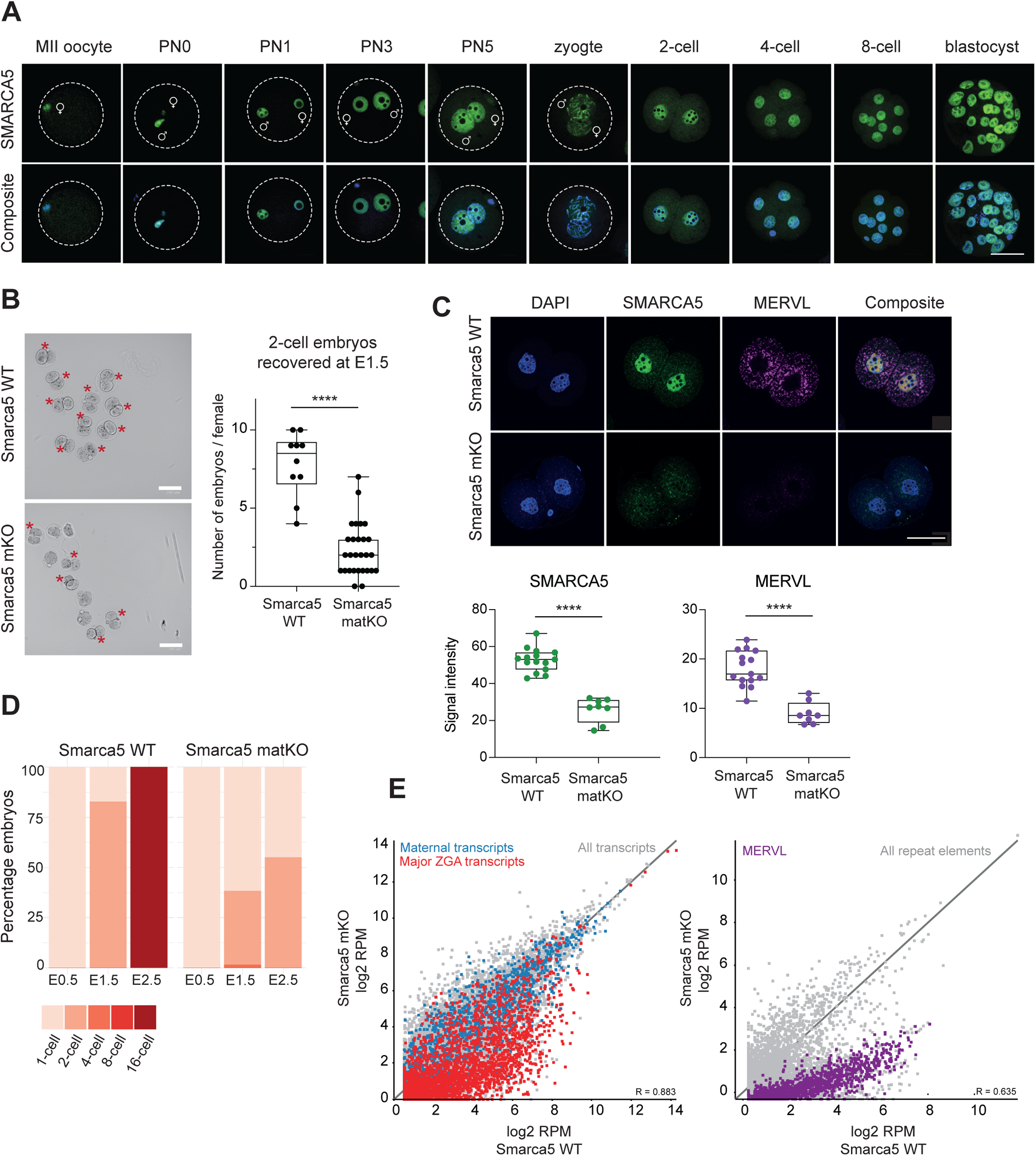
A reduced number of embryos can be recovered from *Smarca5* conditional knock-out females. A. Immunofluorescence images of wild-type CB57L/6 oocytes and preimplantation mouse embryos stained with anti-SMARCA5 antibody (green): Oocyte (MII stage), zygotes (pronuclear stages PN0, PN1, PN3 and PN5, prometaphase 1-cell embryo), 2-cell stage, 4-cell stage, 8-cell stage and blastocyst stage (32-cell). DNA was counterstained with DAPI (blue). Scale bar 50 μm. B. Left: representative brightfield images of wild-type (*Smarca5 WT*) and maternal knockout embryos (*Smarca5 matKO*) collected at E1.5 from wild-type and conditional knockout females (*Smarca5^fl/fl^; Zp3-Cre*) crossed with wild-type male mice. Red asterisks indicate two-cell stage embryos. Right: box plot showing the numbers of two-cell embryos collected at E1.5 from the specified crosses (*Smarca5 WT* n=10 crosses, *Smarca5 matKO* n=28 crosses). Statistical significance was determined by two tailed Mann-Whitney U test. p-value ****P<0.0001, absence of stars (non-significant, ns): p-value>0.05. Scale bar 100 μm. C. Top: Immunofluorescence images of *Smarca5 WT* and *Smarca5 mKO* two-cell embryos stained with α-SMARCA5 (green) and α-MERVL (magenta) antibodies. DNA stained with DAPI (blue), scale bar 50 μm. Bottom: Signal intensity levels for nuclear SMARCA5 and cytoplasmic MERVL in two-cell stage embryos. ****P<0.0001 (Student’s t test). D. Bar plots showing the percentage of embryos that reached each developmental stage, shown by the colours in the legend at the bottom, at the specified time point (E0.5, E1.5 and E2.5). Percentages were calculated based on the total number of zygotes collected at E0.5. E. Scatterplots showing the expression levels (log2 RPM) of maternal transcripts (∼6000, blue), major ZGA genes (3156, red) and MERVL transcripts (732, magenta), in transcriptome data from single embryo NMT-seq libraries, from embryos from one replicate, pooled. Maternal knockout embryos (*Smarca5 matKO*) were compared to wild-type (*Smarca5 WT*) ones.

It was previously reported that oocytes from *Smarca5^fl/fl^; Zp3-Cre* females fail to undergo proper oocyte maturation and do not reach the meiotic stage II (MII) (Zhang et al., 2020). However, when we crossed *Smarca5^fl/fl^; Zp3-Cre* females with wild-type C57BL/6 males, we unexpectedly recovered some two-cell stage embryos at E1.5. The number of embryos was significantly reduced when compared to wild-type *Smarca5*^+/+^ females (Fig. 1B). By immunofluorescence, we observed that the recovered heterozygous embryos (*Smarca5 matKO*) lacked SMARCA5 at the two-cell stage, suggesting that in the absence of maternally deposited SMARCA5, the paternal allele also failed to be expressed at the two-cell stage (Fig. 1C). Moreover, the transposable element MERVL, a prominent ZGA-marker, was not detected in heterozygous embryos from maternal knockout females (Fig. 1C). This shows that major ZGA markers are not properly expressed in absence of maternal SMARCA5. Additionally, heterozygous *Smarca5* matKO embryos did not progress past the two-cell stage *in vitro*, while wild-type embryos at day E2.5 progressed to the morula (16-cell) stage (Fig. 1D). These results suggest that absence of maternal SMARCA5 in oocytes and early embryos is incompatible with ZGA protein marker expression and preimplantation development past the two-cell stage.

To assess if gene expression coupled with epigenetic changes in *Smarca5* maternal knockout embryos were causing the developmental arrest, we performed single-embryo NMT-seq (Clark et al., 2018), which provides simultaneous transcription, DNA methylation and chromatin accessibility data from the same embryo. We collected late two-cell stage embryos, and assessed the expression of major ZGA genes using a gene list that was compiled from several mouse preimplantation RNA-seq data sets (K.-I. Abe et al., 2018; Deng et al., 2014; Liu et al., 2020; Park et al., 2013; Wang et al., 2021; Xue et al., 2013). We concluded that major ZGA genes failed to be upregulated at the two-cell stage in the absence of maternal SMARCA5 (Fig. 1E and Fig. S1C). Similarly, and consistent with the immunofluorescence results, *MERVL* transcripts levels are markedly reduced (Fig. 1E and Fig. S1C). Differential expression analysis between WT and matKO two-cell stage embryos and showed that the majority of differentially expressed genes (DEGs) have lower levels in the absence of SMARCA5 (2030 genes out of 2420) (Fig. S2), and only a few have higher levels. The downregulated DEGs (or genes that failed to be upregulated, Supplementary Table 1) are typically expressed for the first time at the two-cell stage in wild-type embryos, a pattern characteristic of major ZGA genes (Fig. S2A, right boxplots) and enriched for cellular processes like ribosome biogenesis and rRNA processing which are relevant for ZGA (Fig. S2B top). The upregulated DEGs (or genes that fail to be downregulated, Supplementary Table 2) are typically not expressed at high levels in the two-cell stage embryo (Fig. S2B bottom). Altogether, these results demonstrate that in absence of maternal SMARCA5, the expression of major ZGA genes and transposons such as MERVL is impaired in mouse two-cell stage embryos.

### Removal of maternally deposited SMARCA5 in zygotes impairs major ZGA

As the observed developmental failure of *Smarca5 matKO* embryos could be a consequence of the oocyte maturation phenotype previously described in *Smarca5^fl/fl^; Zp3-Cre* females and not its direct involvement in ZGA (Zhang et al., 2020), we decided to remove maternal SMARCA5 protein exclusively after fertilisation in wild-type zygotes that have already undergone normal oocyte development. To achieve this, we adopted Trim-Away, an antibody mediated targeted protein degradation system in early zygotes (Clift et al., 2017) (Fig. 2A).

**Figure 2.**
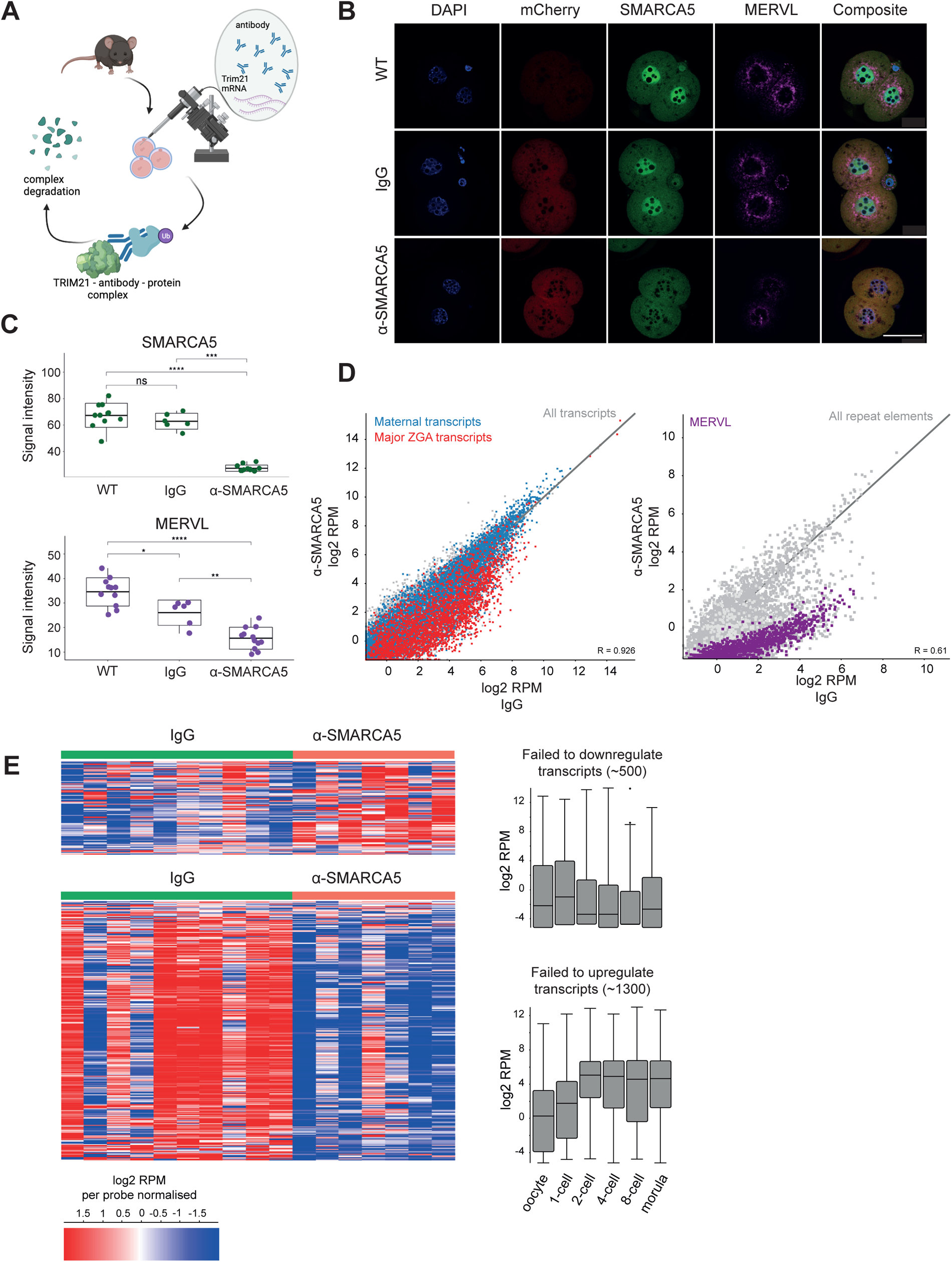
Removing SMARCA5 by Trim-Away impairs major ZGA in two-cell embryos. A. Experimental design of Trim-Away. Zygotes collected from wild-type female mice (C57BL/6J) are microinjected with mCherry-Trim21 mRNA and the anti-SMARCA5 antibody. Once the TRIM21 protein is translated it recognises the Fc-region of the antibody which is already bound to its target, and the whole complex (Trim21-antibody-protein) will be degraded. Following injection, the embryos are cultured for assessment of their developmental potential, immunofluorescence stainings and single embryo NMT-seq. B. Representative immunofluorescence images of two-cell stage embryos from the following three conditions are shown: WT (uninjected), IgG and α-SMARCA5 injected embryos. mCherry-TRIM21 (red), SMARCA5 (green), MERVL (magenta). DNA was stained with DAPI (blue). Scale bar represents 25 μm. C. Box plots with individual data points are showing the signal intensity (raw values, not normalised) for nuclear SMARCA5 and cytoplasmic MERVL protein in each embryo (individual points), replicate 3 is shown. WT (uninjected) n=11, IgG n=8, α-SMARCA5 n=11. Bonferroni adjusted p-values are shown for each comparison, *P<0.05, **P<0.01, ***P<0.001, ****P<0.0001 (Wilcox test). D. Scatterplots showing the expression levels (log2 RPM) of maternal transcripts (∼6000, blue), major ZGA genes (∼3000, red) and MERVL transcripts (∼700, magenta), in the transcriptome data from single embryo NMT-seq libraries generated from two-cell stage SMARCA5 Trim-Away and IgG injected embryos, 28 h post injection, replicate 3. E. Heatmaps (left) showing all DEGs (IgG vs. α-SMARCA5, list compiled from all three experimental replicates) in two-cell stage embryos of replicate 3. Transcripts that failed to downregulate are shown on top, transcripts failed to upregulate at the bottom. The values shown are normalised across datasets (median subtracted from the log2 RPM value for each gene). Box plots (right) illustrate the expression of the corresponding gene sets in wild-type mouse preimplantation embryos (Xue et al. 2013). Boxplots show the median, with the upper and lower extremities of the boxes representing the 25^th^ and 75^th^ percentile of the data.

In order to first validate the efficiency of Trim-Away in our hands, we depleted EG5, a motor protein that is involved in mitotic spindle assembly and has been previously shown to be degradable by Trim-Away (Fig. S3A) in MII oocytes and early zygotes. Following microinjections of anti-EG5 together with the mCherry-Trim21 mRNA, we indeed observed loss of EG5, as well as failure of the formation of the mitotic spindle and absence of alignment of chromosomes at the metaphase plate (Fig. S3B) (Clift et al., 2017). As a second control, we validated the specificity of the anti-SMARCA5 antibody in *Smarca5* knockout mouse embryonic stem cells (*Smarca5*^-/-^ mESCs), and showed absence of any nonspecific immunofluorescence signal. This confirmed that the anti-SMARCA5 antibody we decided to use will most likely not have off-target effects on other proteins in our Trim-Away experiments (Fig. S3C).

To specifically degrade SMARCA5 in early embryos, we injected anti-SMARCA5 (or IgG control) antibody together with mCherry-Trim21 mRNA into early wild-type C57BL/6 zygotes (Fig. 2A). We assessed the degradation efficiency at 7 h (late zygotes) and 28 h (late two-cell stage embryos) post-injection by immunofluorescence. We observed that SMARCA5 protein was not detectable in the late zygotic pronuclei of anti-SMARCA5 injected embryos (Fig. S4A) and was still absent in nuclei of two-cell embryos (Fig. 2B, C, and Fig. S4B). Notably, MERVL protein was significantly reduced in the absence of SMARCA5 at the two-cell stage (Fig. 2B, C and Fig. S4B). In all our experiments, we observed a positive correlation between SMARCA5 levels and the levels of MERVL in Trim-Away embryos (both IgG and anti-SMARCA5 injected) and an inverse correlation between mCherry and SMARCA5 levels (Fig. S4C) which confirmed that the Trim-away system was successfully deployed (i. e. high levels of mCherry tagging TRIM21 protein will rapidly degrade SMARCA5 in those embryos).

The above results suggest that the ZGA associated transposon MERVL fails to be upregulated upon successful degradation of SMARCA5 protein in wild-type embryos. However, while these initial results were similar to the ones observed in *Smarca5* maternal knockout embryos, the developmental potential of anti-SMARCA5 injected embryos seemed only mildly affected, with no significant differences in survival rates between control and SMARCA5 depleted embryos up to blastocyst stage development (Fig. S4D).

To further characterise the molecular consequences of SMARCA5 removal, and assess gene expression, DNA methylation and chromatin accessibility, NMT-seq libraries were generated from late two-cell stage Trim-Away and control embryos 24-28 h post injection. Analysing RNA-seq data from these embryos, we found that major ZGA genes and *MERVL* transcripts failed to be upregulated in the absence of SMARCA5 (Fig. 2D and Fig. S5A, B). Furthermore, DEG analysis identified ∼1300 transcripts that failed to be upregulated upon SMARCA5 depletion (Fig. 2E and Fig. S6, and Supplementary Table 3), including genes characteristic of wild-type two-cell stage ZGA embryos (Fig. 2E, bottom right boxplot). As these two-cell stage and major ZGA specific genes seemed dependent on SMARCA5 in our experiments, they will be further referred to as “SMARCA5-dependent ZGA genes”.

Reassuringly, there was a substantial overlap between genes that failed to be upregulated in SMARCA5 Trim-Away and *Smarca5* matKO embryos. GO term analysis revealed pathways including ‘Ribosome biogenesis’ and ‘RNA processing’, which are characteristic of major ZGA gene expression (Fig. S7A).

In addition, differential expression analysis identified ∼500 genes (compiled from all three replicates) that failed to be downregulated in the absence of SMARCA5 compared to control IgG-injected embryos (Fig. 2E and Fig. S6, and Supplementary Table 4). These transcripts are normally present in wild-type embryos at the zygote stage (Fig. 2E, top right boxplot), indicating that they could be maternal deposits or zygotic transcripts that fail to be removed. Thus, while Trim-Away mediated depletion of SMARCA5 after fertilisation did not recapitulate the developmental phenotype observed in the *Smarca5* maternal knockout embryos, both displayed very similar effects on impaired two-cell stage gene expression. This indicates that SMARCA5 is required for major ZGA transcription in mouse embryos, and not just for oocyte maturation.

### Absence of zygotic SMARCA5 alters accessibility and nucleosome phasing around ZGA gene promoters and enhancers

In order to investigate the molecular details and causes of ZGA impairment (while avoiding potential confounding effects from defective oocyte maturation) we focused on SMARCA5 Trim-Away embryos to assess the effects of its removal at the chromatin level. The main function of SMARCA5 in ISWI complexes is to slide nucleosomes along the DNA (Abdulhay et al., 2021; Barisic et al., 2019; Wiechens et al., 2016), it acts as a nucleosome ruler, and locally and globally regulates chromatin accessibility. We thus hypothesised that changes in ZGA transcription may be caused by altered accessibility or nucleosome positioning around major ZGA genes. To test this, we analysed accessibility data of the generated single-embryo NMT-seq samples from SMARCA5 Trim-Away two-cell embryos.

Notably, we found that transcriptional start sites (TSS) at promoters of SMARCA5-dependent ZGA genes showed lower local accessibility in absence of SMARCA5 as compared to control IgG-injected and wild-type embryos (after background correction; subtraction of median global accessibility) (Fig. 3A, Fig. S8A). TSS at promoters of SMARCA5 independent genes (control promoters), on the other hand, showed no difference between SMARCA5 depleted and control embryos (Fig. 3A, Fig. S8A). SMARCA5 independent gene TSS (controls) are generally less accessible than SMARCA5-dependent ZGA in wild-type embryos at the two-cell stage. In addition, the nucleosome phasing around TSS of major ZGA genes is mildly disturbed in the absence of SMARCA5, as observed in Figure 3A and Figure S8A, based on the different pattern of small peaks following the large accessible TSS peak (see also Fig. S9, non-normalized).

**Figure 3.**
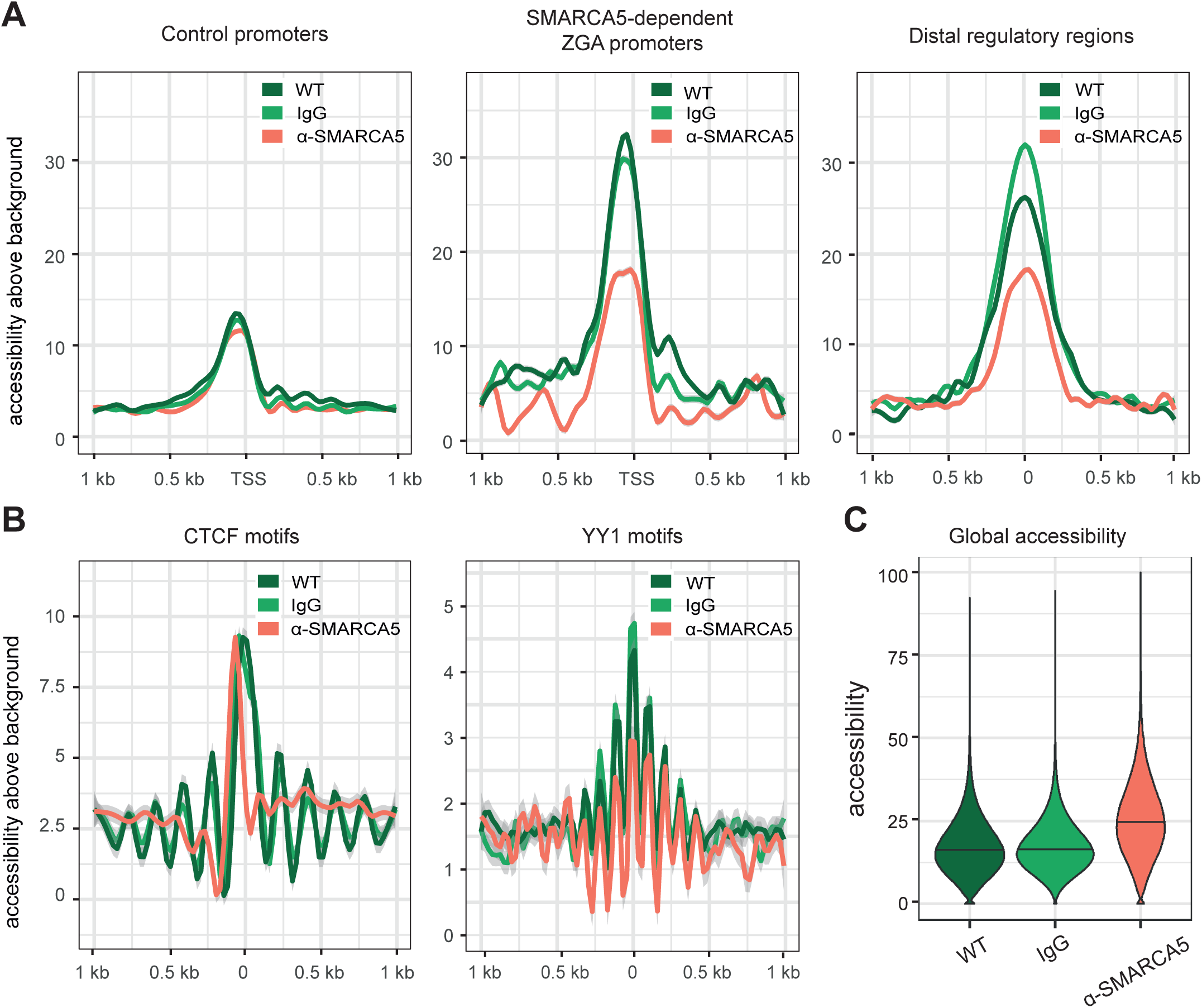
Nucleosome phasing and chromatin accessibility are disturbed in the absence of SMARCA5 at the two-cell stage. A. Nucleosome phasing plots showing accessibility around TSS (+/-1 kb) for control promoters (all promoters except for SMARCA5-dependent gene promoters) and the promoters of SMARCA5-dependent ZGA genes. The plot on the right shows the profile of nucleosomes around the centre of ATAC-seq peaks (+/-1 kb), that was used to define distal regulatory regions (5066 regions). The ATAC-seq peaks were defined in the published dataset from Wu et al 2016. The y axis shows the GpC methylation percentage (readout for accessibility). The data was normalised and shows levels relative to background. Profiles were generated by running 50 bp windows across 2 kb regions. This data shows replicate 3 (IgG n=10 and α-SMARCA5 n=7), the other two replicates are shown in Figure S8. B. Nucleosome phasing plots showing the accessibility and resulting nucleosome profiles around the centre of CTCF (61,068 Homer motifs) and YY1 (57,432 Homer motifs) motifs at the two-cell stage in WT, IgG and α-SMARCA5 embryos. The y axis shows the GpC methylation percentage. The data was normalised, showing the levels relative to background. The profiles were generated by 50 bp running windows across the 2 kb regions. This data is from replicate 3 (IgG n=10 and α-SMARCA5 n=7), the other two replicates are shown in Figure S8. C. Violin plots showing the global GpC methylation percentage, indicative of chromatin accessibility. Accessibility (percentage of methylated GpC sites in each window) was calculated for windows of 200 GpC sites across the genome for both WT embryos or IgG and anti-SMARCA5 injected embryos. This data is from replicate 3 (IgG n=10 and α-SMARCA5 n=7), the other two replicates are shown in Figure S8.

We further examined putative distal regulatory regions that controlled major ZGA genes. These regions were previously identified based on ATAC-seq peaks at the two-cell stage in mice and analysed in SMARCA5 depleted embryos (see Wu et. al., 2016). These distal regulatory elements also showed lower accessibility in the absence of SMARCA5 compared to control embryos, and a slight change in the nucleosome positioning (the small peaks around the centre of the regulatory regions shows a different pattern from the one in control embryos) (Fig. 3A and Fig. S8B). This shows that SMARCA5 is involved in opening of major ZGA gene promoters and distal regulatory regions in mouse two-cell embryos.

In mESC, absence of SMARCA5 has been reported to affect nucleosome phasing around certain transcription factor bindings sites, one of them being CTCF (Barisic et al., 2019; Wiechens et al, 2016). We saw a similar effect in the two-cell stage embryos lacking SMARCA5, with disrupted nucleosome phasing around CTCF binding motifs. This defect seemed specific to CTCF, as the nucleosome positioning around another transcription factor binding motif, YY1, was not affected (Fig. 3B and Fig. S8C, S9B).

In addition to the local changes observed at ZGA gene promoters, putative enhancers and CTCF motifs, the removal of SMARCA5 by Trim-Away seemed to have an effect on global accessibility. Compared to control embryos, the chromatin was on average more accessible in the absence of SMARCA5 across biological replicates (Fig. 3C, and Fig. S8D), and this could also be observed when background normalisation was not applied (Fig. S9).

Importantly, the reduced local accessibility of SMARCA5-dependent ZGA genes, as well as the disruption of nucleosome positioning at ZGA promoters and CTCF motifs could also be observed in *Smarca5* maternal knockout embryos (Fig. S10A, S10B). However, the global increase in accessibility was not observed in these matKO embryos, and we speculate that this is due to the oocyte maturation defect that renders MII oocytes globally less accessible prior to fertilisation (Fig. S10C) (Zhang et al., 2020), and thus a global increase in accessibility would bring them back to similar levels as in control embryos.

Compared to the changes in accessibility in SMARCA5-depleted embryos, the changes in DNA methylation (CpG methylation data generated by scNMT-seq) were less pronounced. We observed a mild increase in global methylation in SMARCA5-depleted embryos, with the majority of the differentially methylated regions showing higher methylation in the absence of SMARCA5 (Fig. S11A). In line with these changes, regions around SMARCA5-dependent ZGA gene promoters and control promoters, were also mildly more methylated in absence of the chromatin remodeler (Fig. S11B).

### Global chromatin organisation changes in the absence of SMARCA5 in two-cell embryos

Following our finding that the chromatin is globally more accessible in the absence of SMARCA5 compared to wild-type two-cell stage embryos, we wanted to investigate other global chromatin changes that might take place at this stage when the chromatin remodeler is not present. The chromatin structure in preimplantation embryos is highly dynamic, with a wide range of changes taking place immediately after fertilisation (Probst & Almouzni, 2011). The number of nucleolus precursor bodies (NPBs) increases in the two-cell stage embryos compared to the zygote, and they are surrounded by “rings” of constitutive heterochromatin, particularly major and minor satellites, marked by HP1-β (Lachner et al., 2001, Probst et al., 2007). These sequences will reorganise into chromocenters at the 8-cell stage, coinciding with the emergence of mature nucleoli (Probst et al., 2007)

To assess if SMARCA5 is required for global chromatin structures developing in embryonic mouse nuclei, we assessed establishment of NPBs in SMARCA5 deficient embryos. Using HP1-β immunofluorescence, we observed structural differences of heterochromatin regions in the nuclei of SMARCA5 deficient embryos as compared to wild-type ones (Fig. 4A). These include a higher number of NPBs in *Smarca5 matKO* embryos, which are, however, smaller in size and contain less HP1-β than NPBs in wildtype embryos (Fig. 4B – 4D, S12A). Similar structural changes in NPB size and number were observed in SMARCA5 Trim-Away embryos. The phenotype was milder (Fig. S12B – S12D), and HP1-β intensity around NPBs was not significantly altered in the latter (Fig. S12E, S12F). Notably, *Smarca5* knockout mouse ESCs also displayed globally altered heterochromatin structures, including HP1-β accumulation in the nucleoli in absence of the remodeler (Fig. S12G). These results suggest that SMARCA5, beyond being required for local accessibility increase at ZGA genes, is also required for global heterochromatin organisation and NPB organisation (see also Fig. 4E).

**Figure 4.**
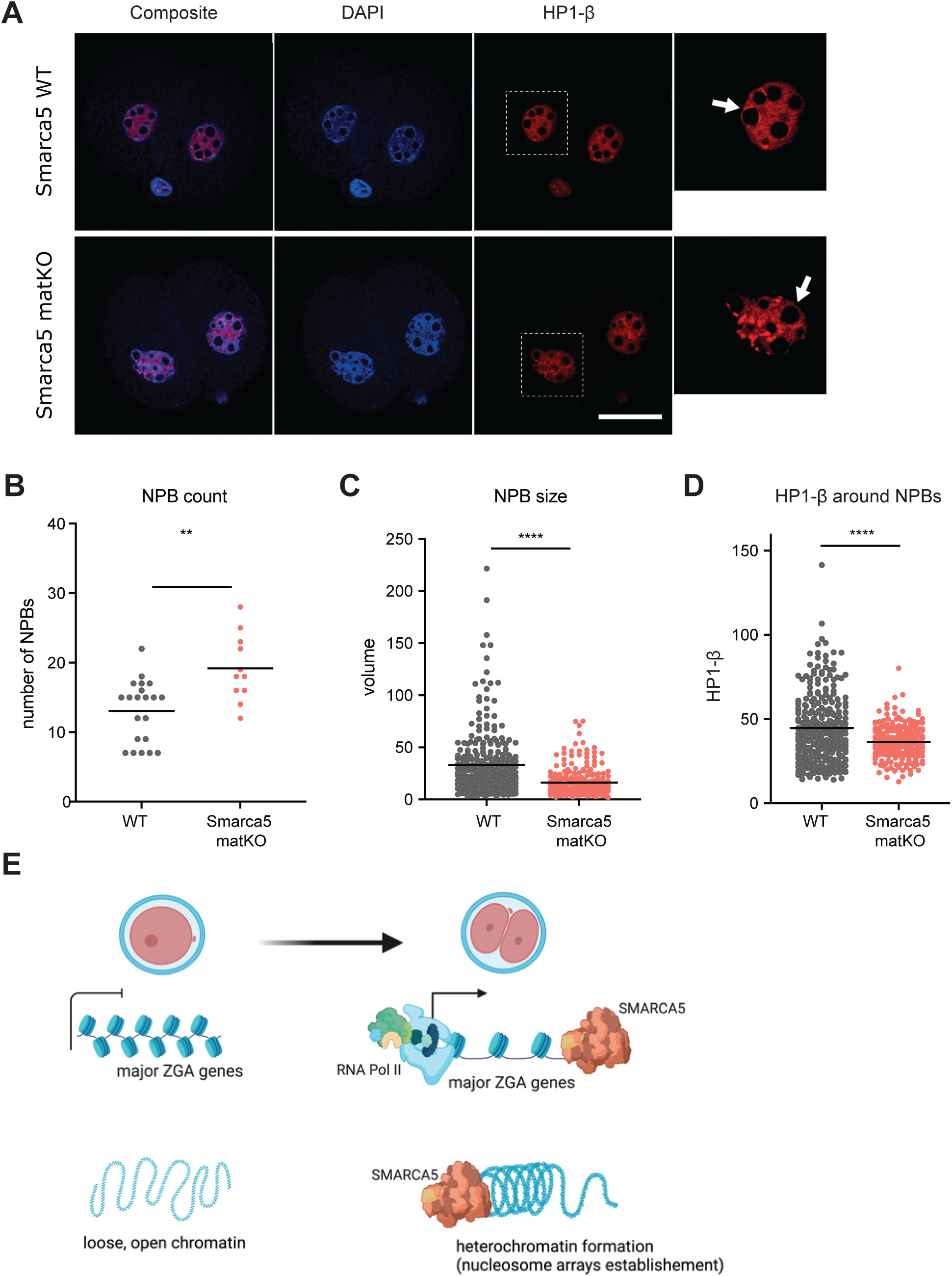
Chromatin structural changes take place at the two-cell stage in the absence of SMARCA5. A. Immunofluorescence images showing representative *Smarca5* wild-type and maternal knockout two-cell embryos stained for HP1-β protein (red). The inserts highlight the nuclei, and the zoomed-in regions are showing the NPBs and the presence (or absence) of their heterochromatic “rings”. Scale bar represents 50 μm. B. Number of nucleolus precursor bodies (NPB count) per *Smarca5* wild-type (WT) and maternal knockout (matKO) 2-cell embryo. Statistical significance was determined by two tailed Mann-Whitney U test. p-value *P<0.05, **P<0.01, ***P<0.001, ****P<0.0001, absence of stars (non-significant, ns): p-value>0.05 C. Volume of NPBs in *Smarca5* wild-type (WT) and maternal knockout (matKO) nuclei as indicated by the longest perimeter. Statistical significance was determined by two tailed Mann-Whitney U test. p-value *P<0.05, **P<0.01, ***P<0.001, ****P<0.0001, ns: p-value>0.05 D. HP1-β signal intensity around NPBs (measured around longest perimeter). Statistical significance was determined by two tailed Mann-Whitney U test. p-value *P<0.05, **P<0.01, ***P<0.001, ****P<0.0001, ns: p-value>0.05 E. Proposed model for the 2 main functions of SMARCA5 in two-cell stage embryos during ZGA. As a regulator of a proportion of ZGA genes (SMARCA5-dependent ZGA genes), SMARCA5 opens up the cromating at their TSS, and also contributes to the nucleosome phasing around this regions, allowing the transcriptional machinery to transcribe these genes. The second role is global, as SMARCA5 seems to be involved in the organisation and reconfiguration of heterochromatic regions in the early embryos.

## Discussion

In this study, we assessed the role of maternally deposited SMARCA5 in mouse ZGA and preimplantation development. We used a maternal conditional knockout mouse model (*Smarca5^fl/fl^; Zp3-Cre* females) to generate *Smarca5* maternal knockout embryos (*Smarca5 matKO*). In addition, we used a targeted protein degradation system to deplete SMARCA5 in wild-type zygotes (SMARCA5 Trim-Away) to show that maternal SMARCA5 is required for mouse ZGA.

*Smarca5* conditional knockout female mice were previously reported to be sterile, due to an oocyte maturation defect (Zhang et al., 2020). However, we recovered a low number of fertilised zygotes and two-cell stage *Smarca5* maternal knockout embryos, which arrested at the two-cell stage, while showing a failure to express major ZGA genes. The observed differences between our study and Zhang et al. could be due to differences in mouse strains or numbers of females used in the study.

To separate the oocyte maturation phenotype from the embryonic phenotype, we adopted the powerful and versatile Trim-Away technology to deplete the chromatin remodeler exclusively after fertilisation in wild-type zygotes. Similar to the maternal knockout embryos, Trim-Away SMARCA5 depleted embryos fail to properly transcribe major ZGA genes. While the major ZGA defect was more severe and consistent among individual *Smarca5* matKO embryos, following Trim-Away we observed a more variable phenotype between replicates and individual embryos throughout our results. This variability is likely caused by differences in the system’s efficiency and reproducibility (introduced by injection efficiency, antibody batch used, and the actual stage of the zygote at the time of injection). Moreover, the variability observed in the Trim-Away embryos can explain the overall lower magnitude of major ZGA gene expression changes, and the reduced impact on the developmental phenotype of these embryos, when compared to their maternal KO counterparts. Using both systems, we concluded that *Smarca5* is a maternal effect gene, as its absence is detrimental for major ZGA and potentially for embryonic development past the two-cell stage.

We further observed that promoters and distal regulatory elements of SMARCA5-dependent ZGA genes failed to open their chromatin in the absence of the chromatin remodeler, suggesting that SMARCA5 specifically increases their accessibility, possibly through local binding. However, based on the global increase in accessibility observed across experiments and conditions in absence of normalisation (Fig. S9), an alternative explanation could be that the SMARCA5-dependent ZGA promoters are as open as in the control embryos, but on the background of a much more globally accessible chromatin landscape that fails to compact in the absence of SMARCA5. Thus, the transcription activation failure of these ZGA genes could be explained either by a direct local role of SMARCA5 at their promoters, or by an indirect global role, which could imply the redistribution of the transcriptional machinery to other regions of the genome (which is overall more accessible in SMARCA5’s absence). A combination of both scenarios might also be possible.

The aberrant enrichment of maternal and minor ZGA transcripts at the two-cell stage in the absence of SMARCA5 is an indication that the embryos fail to efficiently transition from maternal to embryonic control. The degradation of a subset of maternal transcripts is known to be dependent on ZGA (Schultz et al., 2018; Q.-Q. Sha et al., 2019), and thus the failure to activate the embryonic genome could have led to the observed persistence of maternal and zygotic specific transcripts. A similar effect was previously reported upon the removal of other maternal effect genes, such as STELLA/PGC7/DPPA3 (Huang et al., 2017), RNF114 (Zhou et al., 2021), or YAP1 and TEAD4 (Q. Q. Sha et al., 2020).

The embryos in which SMARCA5 was removed by Trim-Away progressed to the blastocyst stage, in contrast to heterozygous embryos from maternal knockouts. However, as the Trim-Away system is transient and the SMARCA5 protein levels could have been restored after the two-cell stage (Israel et al., 2019), we cannot fully exclude that SMARCA5 is transcribed again and contributes to the later stages of preimplantation development. Moreover, it has been previously reported that even if transcription is transiently inhibited at the two-cell stage by a Polymerase II inhibitor, some embryos can still develop to morulae and even blastocyst stages (K.-I. Abe et al., 2018). Therefore, the fact that the embryos in our Trim-Away experiments developed to the morula stage does not entirely exclude the possibility that the ZGA genes transcription at the two-cell stage was affected.

Overall, the above results reveal that SMARCA5 is a key regulator of the activation of the genome in two-cell stage embryos *in vivo*, as was hypothesised following the results from the CRISPR activation (CRISPRa) screen in mESCs *in vitro* (Alda-Catalinas et al., 2019).

We propose a dual role for maternal SMARCA5 in mouse preimplantation development and ZGA (Fig. 4E). Our results reveal that SMARCA5 is required to position nucleosomes locally at ZGA promoters, regulatory elements (MERVL) and transcription factor binding sites (CTCF). In this context SMARCA5 opens the chromatin of ZGA associated genes and regulatory elements and contributes to initiation of their transcription.

We also observed that SMARCA5 contributes to *de novo* heterochromatin formation during early development, its absence leading to global structural changes, as witnessed by the global increase in chromatin accessibility and the aberrant HP1-β distribution at the two-cell stage.

## Author contributions

Conceptualization: W.R., J.T.S., O.K., M.E.M.; Methodology: O.K., F.S., D.D., M.E.M, C.A.C; Formal analysis: O.K., F.S., H.O., C.K.; Investigation: O.K., F.S., C.K., J.T.S.; Data curation: O.K., F.S., C.K.; Writing – original draft: O.K., J.T.S.; Writing – review & editing: O.K., J.T.S., W.R., F.S.; Visualization: O.K., F.S., J.T.S.; Supervision: W.R., J.T.S.; Funding acquisition: W.R.

## Acknowledgements

We thank all members of the Reik laboratory for their helpful discussions. We thank Wendy Dean for helpful advice on mouse genetics, breeding and colony management. We thank all staff in the Babraham Biological Support Unit (BSU), Felix Krueger, Laura Biggins, and Simon Andrews in the Bioinformatics facility, Simon Walker in the Imaging Facility and Nicole Forrester and Paula Kokko-Gonzales in the Sequencing Facility at Babraham Institute for their support.

## Data availability

RNA-seq and NMT-seq data have been deposited in GEO under accession number <u>GSE248181</u>.

## Methods

### Embryo collection and culture

All mouse embryos were collected from natural matings or IVF (*in vitro* fertilisations) between the appropriate genotypes or wild-type animals according to standard procedures (Hogan, 1994) at different time points, depending on the desired stage.

MII wild-type oocytes were collected from C57Bl/6 females in oestrus. To obtain zygotes and two-cell embryos, at E0.5 and E1.5 respectively, we set up the C57Bl/6 females in matings with C57Bl/6 males. *Smarca5* maternal knockout embryos (*Smarca5* matKO, heterozygous for *Smarca5*) and MII oocytes were derived from *Smarca5^fl/fl^; Zp3-Cre* females, mated with C57Bl/6 wild-type males to obtain the embryos.

For Trim-Away experiments, C57Bl/6 wild-type females were superovulated by hormonal injections with 5 IU pregnant mare serum gonadotropin (PMSG) and 5 IU human chorionic gonadotropin (hCG) 48 hours after PMSG, then set up in matings with C57Bl/6 wild-type males. For *in vitro* culture, embryos and oocytes were collected in M2 media (Sigma-Aldrich, MR-015P-5F) containing hyaluronidase (Sigma, H2126) and further washed in M2 drops to remove the cumulus cells (for oocytes and 1 cell stage only). All embryos were cultured in KSOM media made in-house, covered with mineral oil (Sigma, ES-005-C), at 37°C, 5% CO2. For the assessment of the developmental potential, Trim-Away embryos were cultured in ORIGIO® Sequential Cleav™ (CopperSurgical, 83040010) up to the 8 cell stage, and in ORIGIO® Sequential Blast™ (CopperSurgical, 83050010) up to the blastocyst stage.

### *In vitro* fertilisation

*In vitro* fertilisation (IVF) was used to obtain the embryos used for timed immunofluorescence experiments, to have better temporal control over the early zygotic stages. Four- to five-week-old C57Bl/6 females were superovulated and MII oocytes collected in fertilisation media (Human Tubal Fluid (HTF, MR-070, Sigma) containing 4 mg/ml BSA and 25mM L-Glutathione 1 (GSH)). The sperm collected from one epididymis was released in HTF, and then incubated for 30 min at 37°C, 5% CO2. The sperm with the highest mobility was then collected from the edge of the drop and placed in the fertilisation media drop together with the oocytes. They were incubated for 5 hours at 37°C, 5% CO2, with batches of fertilized embryos collected every hour within that time window, washed in PBS and fixed for further experiments.

### Trim-Away

mCherry-Trim21 mRNA cloning and *in vitro* transcription: The plasmid pGEMHE-mCherry-mTrim21 was a kind gift from Melina Schuh (Adgene plasmid #105522; http://n2t.net.addgene:105522; RRID:Addgene 105522). The mCherry-Trim21 fragment was PCR amplified (Phusion high fidelity DNA polymerase, MO530S) to introduce ClaI restriction sites (fwd TAATTATCGATTATAATGGTGAGCAAGGGCGAGGA, rev TATTAATCGATCCGCTCACATCTTTAGTGGACAGA) The PCR product was purified (NEB Monarch PCR and DNA purification kit, T1030S), digested with ClaI (NEB, R0197S) for 2 hours, 37°C and cloned into the ClaI linearized transcription efficient plasmid pCS2+ (a kind gift from Anna Philpott’s lab). The resulting plasmid was transformed into and amplified in competent DH5α cells (Invitrogen, Cat no. 18263-012), purified (QIAprep Spin Miniprep kit, Cat no: 27104) the sequence confirmed with Sanger sequencing (Sp6 promoter primer and T3 primer). *In vitro* transcription was done according to the SP6 mMESSAGE mMACHINE kit manual (ThermoFisher, AM1340), followed by 15 minutes of DNase treatment, and polyA tailing for 1 h, 37°C (ThermoFisher, AM1350). The mRNA was purified (RNeasy Mini kit, QIAGEN, Cat no: 74104), eluted in 20 μl of nuclease-free water, aliquoted and stored at -80°C. The mRNA translation efficiency was tested by transfection into mESCs (Lipofectamine MessengerMAX Transfection Reagent, ThermoFisher, LMRNA003) and monitoring of mCherry expression after 6 and 22 h (ZOE fluorescent cell imager, Bio-Rad, 1450031).

Antibody purification and concentration: The anti-Smarca5 (Abcam, ab72499), and rabbit IgG (Merk-Millipore, 12-370) antibodies were purified using the Amicon Ultra-0.5 Centrifugal Filter Units (Merck, UFC510024) as follows: 500 μl antibody was added to the column, centrifuged for 10 min at 4° C, 14,000 g, washed three times with 480 μl PBS (10 min at 4° C, 14,000 g). The column was then reversed in a new collection tube, centrifuged for 2 min, 1,000 g at 4° C (see also manufacturer’s protocol) and 20 μl of purified antibody in PBS were recovered. The concentration was measured using Nanodrop2000 (1 Abs=1mg/ml; since the extinction coefficient for IgG is 1.4 (1 Abs=0.714 mg/ml), the value was multiplied by 0.714 to obtain the final antibody concentration (Clift et al., 2018). The purified antibodies were aliquoted, snap-frozen in liquid-nitrogen and stored at -80°C.

Trim-Away mix preparation: Just before microinjection, the Trim-Away mix was freshly prepared on ice; tubes and tips were UV-treated 1 hour prior, and the bench was cleaned with RNaseZAP (ThermoFisher, AM9782). The final injection mix contained: mCherry-Trim21 mRNA (200 ng/μl), IgG or anti-Smarca5 antibody (0.5-1 mg/ml), NP-40 (0.05%), and PBS to 25 μl. The solution was kept on ice until it was loaded into the microinjection needles.

Microinjections: Early zygotes were collected from superovulated four- to 4-5 week-old C57Bl/6 females crossed with C57Bl/6 males. The zygotes were cleaned in M2 medium (Sigma-Aldrich, MR-015P-5F) with added hyaluronidase (Sigma, H2126) and kept in culture in KSOM until they were injected. M2 medium was used as handling media during injections. The embryos were injected using a Nikon eclipse TE2000-U inverted microscope and Narishige micromanipulators, and were moved back into KSOM and cultured until the required time point.

### Mouse embryonic stem cell culture

Wild-type (E14) and knockout (*Smarca5*-/-) mouse embryonic stem cells (mESC) were cultured in serum/LIF conditions: DMEM (Gibco, 11995-040), 15% fetal bovine serum (FBS), 1 U/mL penicillin – 1 mg/mL streptomycin (Gibco, 15140-122), 0.1 mM nonessential amino acids (Gibco, 11140-050), 4 mM GlutaMAX (Gibco, 35050-061), 50 μM β-mercaptoethanol (Gibco, 31350-010), and 103 U/mL LIF (Stem Cell Institute, Cambridge). The cells were maintained on culture plates coated with 0.1% gelatine, at 37°C with 5% CO2 humidified atmosphere, and passaged with Trypsin EDTA (Thermo Fisher Scientific, 25200056). *Smarca5* knockout cells were previously described in Barisic et al., 2019.

### Immunofluorescence

*Oocytes and embryos* were washed in PBS, fixed for 15 min in 4% paraformaldehyde (PFA) in PBS, and stored in 0.05% sodium azide (NaAZ) in 0.05% Tween20 in PBS (PBT) at 4°C. For staining, they were permeabilized using 0.5% TritonX-100 in PBS for 1 h, then blocked for 1 h in 1% BSA in PBT (BS). Primary antibodies (see Antibody list below), were diluted in BS, incubated with the sample for 1 h, and embryos then washed for 1 h in BS. Secondary antibodies diluted 1:1000 in BS were incubated for 45 min RT, and washed for 1 hour in PBT. Secondary antibodies used were Alexa Fluor (Invitrogen) anti-mouse, anti-goat, anti-rat or anti-rabbit IgG conjugated with different fluorophores, depending on the experiments. DNA was counterstained with 5 μg/ml DAPI for 10 min. All incubations were performed at room temperature. The embryos were mounted in fibrin clots on glass slides (Hodges et al., 2002), using Prolong Gold Antifade mounting media (ThermoFisher, P36934). Single optical sections or Z-stacks were captured with a Zeiss LSM780 microscope (63x oil-immersion objective). Images were pseudo-coloured and corrected for contrast and brightness within the recommendations for scientific data. Fluorescence semi-quantification analysis was performed with Volocity 6.3 software (Improvision) and ImageJ. The nuclei of each blastomere in the two-cell embryos were measured separately. The plots were generated using RStudio.

*Mouse ESCs* from 10 cm culture plates were harvested and disassociated using Trypsin EDTA (Thermo Fisher Scientific, 25200056), washed twice in PBS (cells pelleted by centrifugation 300 rpm, 3 min), and resuspended in 2 ml 2% PFA (Polysciences, Inc., 18814). After 30 min incubation at room temperature (RT), the PFA was removed by centrifugation and the cells resuspended in PBS containing 0.05% NaAZ for storage at 4°C until further use. For staining, the cells were concentrated and cytospun onto glass slides (300 rpm, 3 min), and permeabilised with 0.5% TritonX-100 in PBS for 30 min at RT, in a humidified chamber. This was followed by blocking for 30-60 min at RT in BS. Primary antibodies were diluted in BS (see Antibody list below), and incubated with the samples for 1 h RT, followed by three 10 minute washes in PBT (0.05% Tween20/PBS). Secondary antibodies were diluted 1:1000 in BS and incubated for 30 min RT, then washed for at least one hour in PBT at RT. Secondary antibodies used were Alexa Fluor (Invitrogen) conjugated anti-mouse, anti-goat, anti-rat or anti-rabbit IgGs conjugated with different fluorophores, depending on the experiments. DNA was counterstained with 5 μg/ml DAPI in PBS solution for 10 min RT, and coverslips were mounted in Prolong Gold Antifade mounting media (ThermoFisher, P36934).

### Antibody direct labelling and immunofluorescence

In order to check whether the system worked and the protein was depleted, the embryos were fixed in 4% PFA at approximately 7 and 24 hours after injection, followed by immunofluorescence staining, however, as the embryos already contained an injected antibody against the protein of interest, direct labelling of antibodies had to be used in some cases. The primary antibodies were directly labelled with one of the AF488, AF568 or AF647 labelling dyes, following the Zenon™ Alexa Fluor™ Rabbit Labelling Kit instructions (Molecular Probes A20181, A20184 and A20186). Primary antibody incubation was carried out for 1 hour at RT, in the dark, followed by three washes in PBT (15 min each), and a short fixation step for 15 min in 2% PFA at RT. After a 10 minutes PBS wash, the DNA was stained with DAPI (5 μg/ml) for 10 minutes, then the embryos were mounted in fibrin clots (Hodges et al., 2002) with Prolong Gold Antifade mounting media (ThermoFisher, P36934).

### NPB analysis

A pipeline for image analysis was developed in Fiji and R. First, the StarDist neural network (Schmidt 2018, Chamier 2021) was trained to recognise NPBs, using a training dataset of paired images of confocal z-slices and their ground truth NPB segmentation masks. The newly trained StarDist_NPB network was then implemented in a Fiji script to enable segmentation of NPB masks across the z-slices of the confocal image stack, which were subsequently linked in 3D to form 3D NPB objects using a custom TrackMate (Tinevez 2017) script. The volume of NPBs objects was analysed using the MorpholibJ plugin (Legland 2016). Also extracted was the HP1β signal intensity along the periphery of each NPB in each optical slice it appeared in.

### Single-embryo RNA-seq

Smarca5 Trim-Away and control embryos were generated as indicated. The zonae pellucidae were removed using Tyrode’s solution (Sigma-Aldrich, T1788) and individual embryos were placed in 2.5 µl methyltransferase reaction mixture, according to the published protocol (Clark et al., 2018). mRNA was captured using Smart-seq2 oligo-dT pre-annealed to magnetic beads (MyOne C1, Invitrogen). The lysate containing the gDNA was transferred to a separate PCR plate and the beads were washed three times in 15 ml FSS buffer (Superscript II, Invitrogen), 10 mM DTT, 0.005% Tween 20 (Sigma-Aldrich) and 0.5 U/ml of RNAsin (Promega). The beads were then resuspended in 10 ml of reverse transcriptase mastermix [100 U SuperScript II (Invitrogen), 10 U RNAsin (Promega), 1× Superscript II First-Strand Buffer, 2.5 mM DTT (Invitrogen), 1 M Betaine (Sigma-Aldrich), 9 mM MgCl2 (Invitrogen), 1 mM Template-Switching Oligo (Exiqon), 1 mM dNTP mix (Roche)] and incubated on a thermocycler for 60 min at 42°C, followed by 30 min at 50°C and 10 min at 60°C. PCR was then performed by adding 11 ml of 2× KAPA HiFi HotStart ReadyMix and 1 ml of 2 mM ISPCR primer, and cycling as follows: 98°C for 3 min, 15 cycles of 98°C for 15 s, 67°C for 20 s, 72°C for 6 min and finally 72°C for 5 min. cDNA was purified using a 1:1 volumetric ratio of Ampure Beads (Beckman Coulter) and eluted in 20 ml of water. Libraries were prepared from 100 to 400 pg of cDNA using Nextera XT Kit (Illumina), as per the manufacturer’s instructions but with one-fifth volumes for each sample. Libraries were sequenced on an Illumina NextSeq500 MidOutput 75 bp paired-end reads per embryo. Data available at GEO under accession number GSE184763.

### RNA-seq analysis

Libraries were trimmed using Trim Galore (v0.6.5, Cutadapt v2.3) and mapped to the mouse GRCm38 genome assembly using HISAT2 (v2.1.0, --no-softclip) and filtered to have MAPQ scores of 20 and above. Data were quantified using the RNA-seq quantitation pipeline in SeqMonk (https://www.bioinformatics.babraham.ac.uk/projects/seqmonk/), quantifying reads from both strands, using mRNA probes. All the analysis, specified in each figure legend, was carried in SeqMonk, and only the violin plots were generated in RStudio. The differentially expressed genes (DEGs) were identified using the DESeq2 pipeline in Seqmonk (p<0.05). The GO term analysis figures were generated with the online tool ShinyGO 0.76 (http://bioinformatics.sdstate.edu/go/). The major ZGA and maternal deposited gene lists used in the analysis were generated using the publicly available dataset (GSE44183) from Xue et al. (2013). The 2C-like gene list was from Eckersley-Maslin et al. (2016).

### Single embryo Nucleosome, Methylation and Transcription sequencing (NMT-seq)

Embryo collection: Mouse MII oocytes, 1 and 2 cell embryos from Trim-Away experiments or conditional knock-out females were washed in PBS, the zona pellucida was removed using Tyrode’s solution (Sigma-Aldrich, T1788), and embryos were placed in 2.5 μl GpC methyltransferase (M.CviPI, M0227, NEB) reaction buffer at 37°C for 30 min, according to the published protocol (Clark et al., 2018). 5 μl RLT (79216, Qiagen) was added, samples were snap-frozen and stored at -80°C. The two blastomeres of 2 cell stage embryos were processed together as one sample in all experiments.

NMT-seq library generation and sequencing: Library generation (96-well) was done as previously described (https://www.protocols.io/view/scnmt-seq-kxygxmwwwl8j/v3, Clark et al., 2018). Briefly, the RNA and gDNA were physically separated using oligo(d)T-conjugated magnetic bead mRNA capture (MyOne C1, Invitrogen) and transfer of the gDNA containing lysate to a separate PCR plate.

scRNA-seq libraries: Oligo-d(T) beads with RNA were washed three times [15 μl FSS buffer (Superscript II, Invitrogen), 10 mM DTT, 0.005% Tween 20 (Sigma-Aldrich) and 0.5 U μl-1 RNAsin (Promega)]. The mRNA was reverse transcribed in 10 μl reverse transcriptase mastermix [100 USuperScript II (Invitrogen), 10 U μl-1 RNAsin (Promega), 1Å∼ Superscript II First-Strand Buffer, 2.5 mM DTT (Invitrogen), 1 M Betaine (Sigma-Aldrich), 9 mM MgCl2 (Invitrogen), 1 mM Template-Switching Oligos (Exiqon) (Picelli et al., 2013), 1 mM dNTP mix (Roche)] for 60 min at 42°C, followed by 30 min at 50°C and 10 min at 60°C. After the addition of 11 μl of 2Å∼ KAPA HiFi HotStart ReadyMix (7958927001, Roche) and 1 μl of 2 mM ISPCR primers (Picelli et al. 2013), the samples were amplified under the following conditions: 98°C for 3 min, 15 cycles of 98°C for 15 s, 67°C for 20 s, 72°C for 6 min and finally 72°C for 5 min. The resulting DNA was purified (Ampure Beads, Beckman Coulter) and eluted in 20 μl nuclease-free water. Libraries were prepared from 100-400 pg of cDNA using Nextera XT Kit (Illumina), as per the manufacturer’s instructions but with one-fifth volumes for each sample.

scNOME-seq libraries: The cell lysate containing the genomic DNA, was purified with a 0.8:1 volumetric ratio of Ampure XP Beads (Beckman Coulter), washed twice in 80% EtOH and eluted into 10 μl of nuclease-free water. Bisulfite conversion was carried out using EZ Methylation Direct MagBead kit (Zymo) using half of the volumes recommended in the manufacturers’ protocol, for 3 h, at 64°C, followed by desulphonation and purification. Converted DNA was eluted in 40 μl of first strand synthesis mastermix [1 Å∼ Blue Buffer (Enzymatics), 0.4 mM dNTP mix (Roche), 0.4μM 6NF oligo (IDT)]. On the thermocycler, the DNA was heated to 65 °C for 3 min and immediately cooled on ice, to allow the addition of 50 U of Klenow exo-(Enzymatics), and from 4°C slowly ramping up to 37 °C, incubated for 30 min. This was repeated 4 more times, but following the initial incubation at 65°C, 2.5 μl of reaction mixture prepared in advance was added at every round: 1Å∼ Blue buffer, 0.25 mM dNTPs, 10 mM 6NF oligo and 50 U Klenow exo-(Enzymatics). After the first-strand synthesis, 40 U of Exonuclease I (NEB, M0568) was added and the reaction was incubated at 37 °C for 1h, and purified using a 0.75:1 ratio of AMPure XP beads. A mix containing 50 μl of second strand mastermix [1Å∼ Blue Buffer (Enzymatics), 0.4 mM dNTP mix (Roche), 0.4 μM 6NF oligo (IDT)] was used to resuspend the purified products, which were then heated at 98 °C for 2 min and cooled on ice. 50 U of Klenow exo-(Enzymatics) was added, and the plate was incubated on a thermocycler, slowly ramping up the temperature from 4°C until 37°C, where it was kept for 90 min. The obtained second strand products were purified using a 0.8:1 ratio of AMPure XP beads. The PCR mastermix (50 μl) containing 1Å∼ KAPA HiFi Readymix, 0.2 μM PE1.0 primer, 0.2 μM iTAG index primer, was used to elute the products. In order to amplify the final scBS-seq library, the reaction was cycled for 2 min at 95°C, followed by 14 cycles of 80 sec at 94°C, 30 sec at 65°C, 30 sec at 72°C, and extended for 5 min at 72°C. The samples were then pooled (5 μl of each sample), and the PCR reaction components were removed by a sequential purification using 0.7:1 and 0.8:1 volumetric ratio of AMPure XP beads to reaction volume.

The quality and concentration of all final libraries was checked using a Bioanalyzer device (Agilent). The multiplexed scRNA-seq libraries were sequenced at 75 bp paired-end on the Illumina NextSeq500 MidOutput and the multiplexed single-cell bisulfite and NOME-seq libraries were sequenced at 100 bp paired-end on the Illumina HiSeq2500 Rapid Run platform.

### scRNA-seq (from single embryo NMT-seq) data processing and analysis

Raw FastQ files were trimmed using Trim Galore (v0.6.5, Cutadapt v2.3) and mapped to the mouse GRCm38 genome assembly using HISAT2 (v2.1.0) (D. Kim et al., 2015), with the options –dta –no-softclip –no-mixed –no-discordant. Only the reads that had MAPQ scores of 20 and above were retained.

The mapped data was imported into SeqMonk (www.bioinformatics.babraham.ac.uk/projects/seqmonk/) and the outlier samples (> 15% mitochondrial reads, < 25% genes) discarded . Gene expression was quantified using the RNA sequencing quantitation pipeline (mRNA probes, correcting for the total number of sequences in the dataset, and generating log transformed values (log 2 RPM)). Differential expression analysis between wild-type and maternal KO embryos or Trim-Away SMARCA5 depleted embryos was performed using DESeq2 (with multiple test correction, p <0.05) and Intensity Difference (with multiple test correction, p <0.05), followed by the intersection of the two.

Repeat elements expression levels (MERVL, LINE1) were measured by quantitating the individual instances of these repeats in the genome, and only uniquely mapped instances were counted (Read count quantitation pipeline, normalised to the total number of sequences in the dataset, log transformed (log2 RPM)). For data visualisation, plots were either directly generated in SeqMonk, or the data was exported and the final plots generated in R.

### NOME-seq (from single embryo NMT-seq) data processing and analysis

FastQ files were trimmed with Trim Galore (v0.6.5, default parameters) and mapped to the mouse GRCm38 genome assembly using Bismark (v0.23.1) (Krueger & Andrews, 2011), with the additional --NOMe option in the coverage2cytosine script and –CX for the methylation extraction and coverage2cytosine processes. This produced separate report files: CpG files containing only A–C–G and T–C–G positions and GpC files containing only G–C–A, G–C–C and G–C–T positions. On average, more than 33% of C in a CpG context and more than 4.5% of C in a GpC context were methylated, as we would expect from this type of data for quality control (Clark et al., 2018). These CpG and GpC report files were imported into SeqMonk for data analysis.

DNA methylation: For DNA methylation analysis, data was quantitated using the bisulfite methylation over features SeqMonk pipeline, over non-overlapping windows containing 100 CpGs, spanning the entire genome, calculating the percentage methylation for each base (C) and averages these for each measured window. The windows were measured only if they had at least 10 observations (10 CpG sites out of 100, that were covered by at least 1 read). Because of the data sparsity, the analysis was carried out in pseudo-bulk, merging the data from individual embryos belonging to the same replicate and condition. On average 140,000 windows were measured for each pseudo-bulked sample. The same 100 CpG windows were used to assess the methylation status of different genomic features overlapping these windows (e.g. promoters and CpG islands). Differential methylation analysis was done in SeqMonk using the proportion based statistical test Chi-Square (with multiple test corrections, p< 0.05) together with EdgeR (p<0.05).

Accessibility data: Since the levels of C methylation in the GpC context is the read-out for accessibility, the analysis is similar to the one carried out on conventional CpG methylation. GpC data was quantitated using the Bisulfite Methylation over features pipeline, over 200 GpC (global accessibility) or 300 GpC (differentially accessible regions) containing windows, spanning the entire genome. The windows containing less than 10 observations (covered GpC sites, by at least 1 read) per window were filtered out, and coverage outliers (> 100 RPM) in any sample were excluded. Due to data sparsity, datasets from individual embryos were merged according to their replicate and condition, and the rest of the analysis was carried out in pseudo-bulk. For the evaluation of accessibility at specific regions, only the 200 GpC windows that overlapped the genomic features of interest (promoters, distal regulatory regions) were considered.

For the nucleosome occupancy analysis at promoters, 50 bp probes (windows) were generated across regions 1 kb upstream and downstream of all gene start sites and quantitated using the bisulfite methylation pipeline (with minimum 1 GpC covered by at least 1 read).

For nucleosome occupancy at distal regulatory regions, the 50 bp probes were made across the 3 kb surrounding the centre of promoter distal ATAC-seq peaks, previously defined from a published mouse preimplantation ATAC-seq dataset (J. Wu et al., 2016a). For phasing around the CTCF motifs (HOMER motifs), the 50 bp probes were made across 2 kb from the centre of the motif.

### Other software for figure generation

All the plots in this manuscript were generated in RStudio and SeqMonk (https://www.bioinformatics.babraham.ac.uk/projects/seqmonk/). The schematic figures were generated with BioRender (https://biorender.com/) and modified in Adobe Illustrator.

### Quantification and statistical analysis

#### Statistics and reproducibility

Statistical tests, sample sizes and definitions of error bars of each experiment are indicated in the figure legends and were calculated using Graphpad Prism (version 9.2.0). For all tests, p values were presented as *p<0.05, **p<0.01, and ***p<0.001.

#### Primary antibodies

rabbit anti-EG5 (Merck-Millipore, HPA010568)

rabbit anti-MERVL (Huabio, R1501-2, IF: 1:100)

rabbit anti-SMARCA5 (abcam, ab72499, IF: 1:200)

rabbit anti-mCHERRY (abcam, ab167453, IF: 1:200)

rat anti-HP1β (abcam, ab10811, IF: 1:300)

mouse anti-pan-histone (Merck-Millipore, mab3422, IF: 1:400)

#### Secondary antibodies

anti-rabbit AlexaFluor-conjugated 647 (Molecular Probes, A31573, IF: 1:1000)

anti-mouse AlexaFluor-conjugated 568 (Molecular Probes, A10037, IF: 1:1000)

anti-goat AlexaFluor-conjugated 488 (Molecular Probes, A11055, IF: 1:1000)

