## Supplemental Figures for "Maternal SMARCA5 is required for major ZGA in mouse embryos"

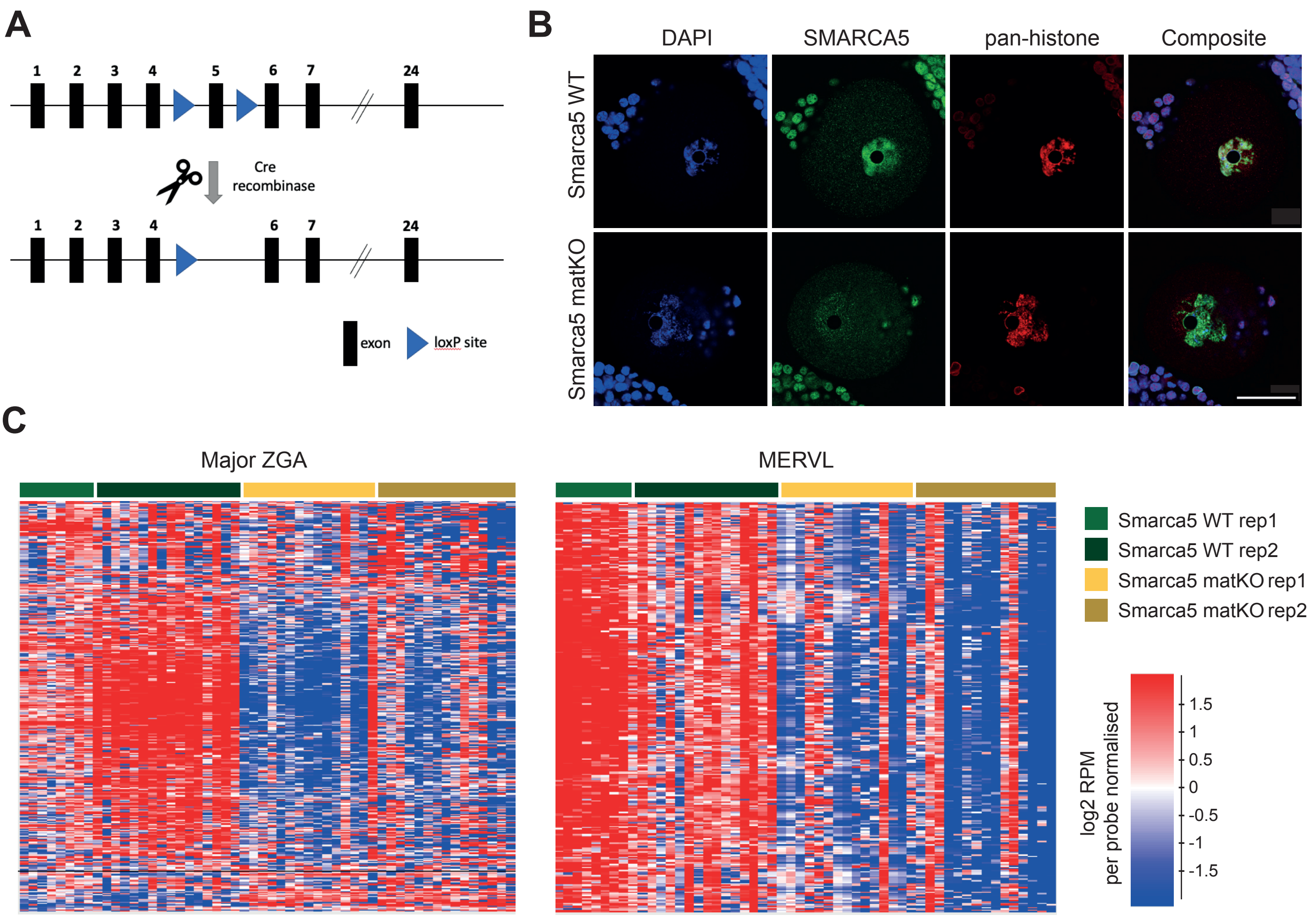


**Supplemental Figure 1**

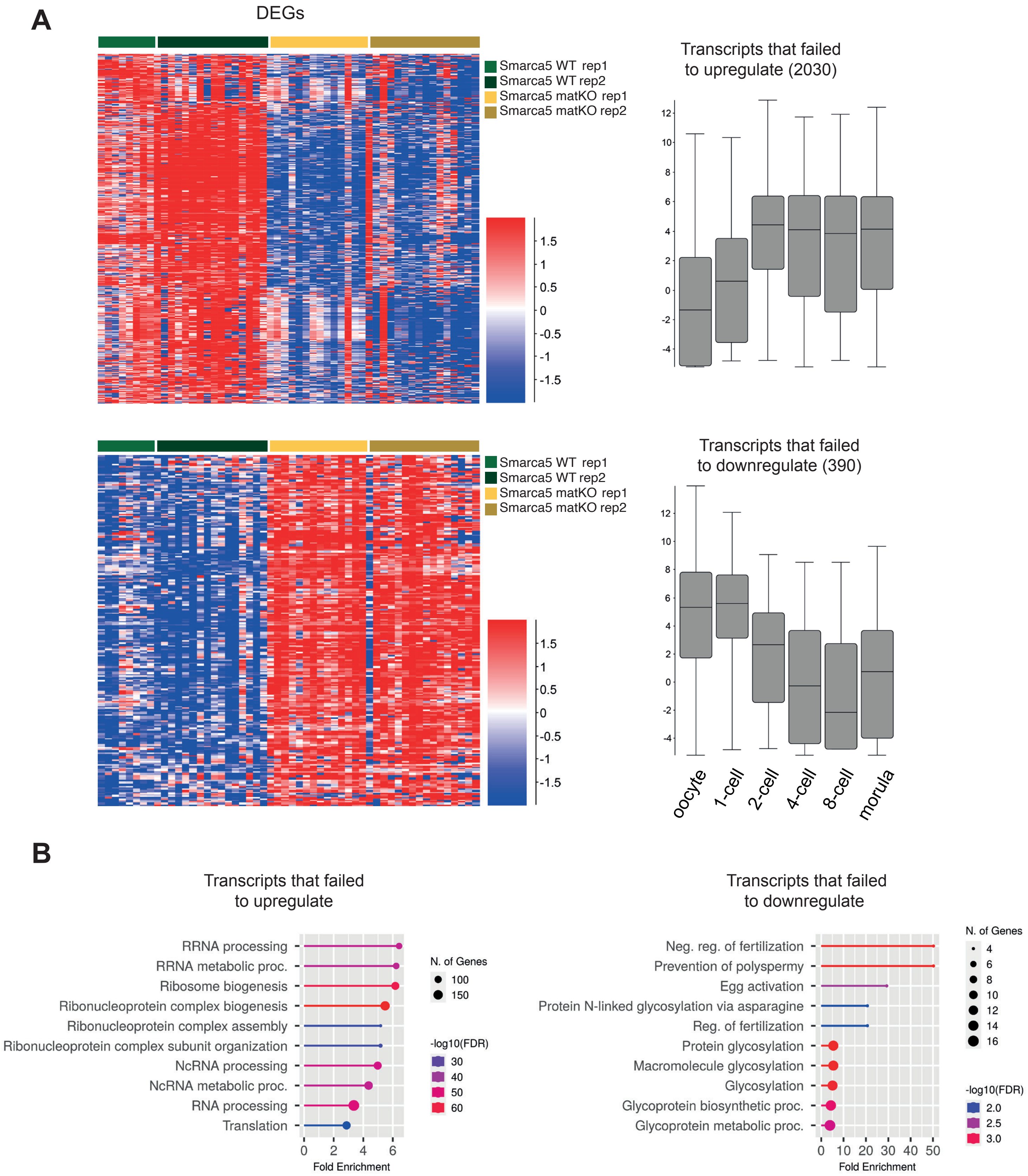


**Supplemental Figure 2**

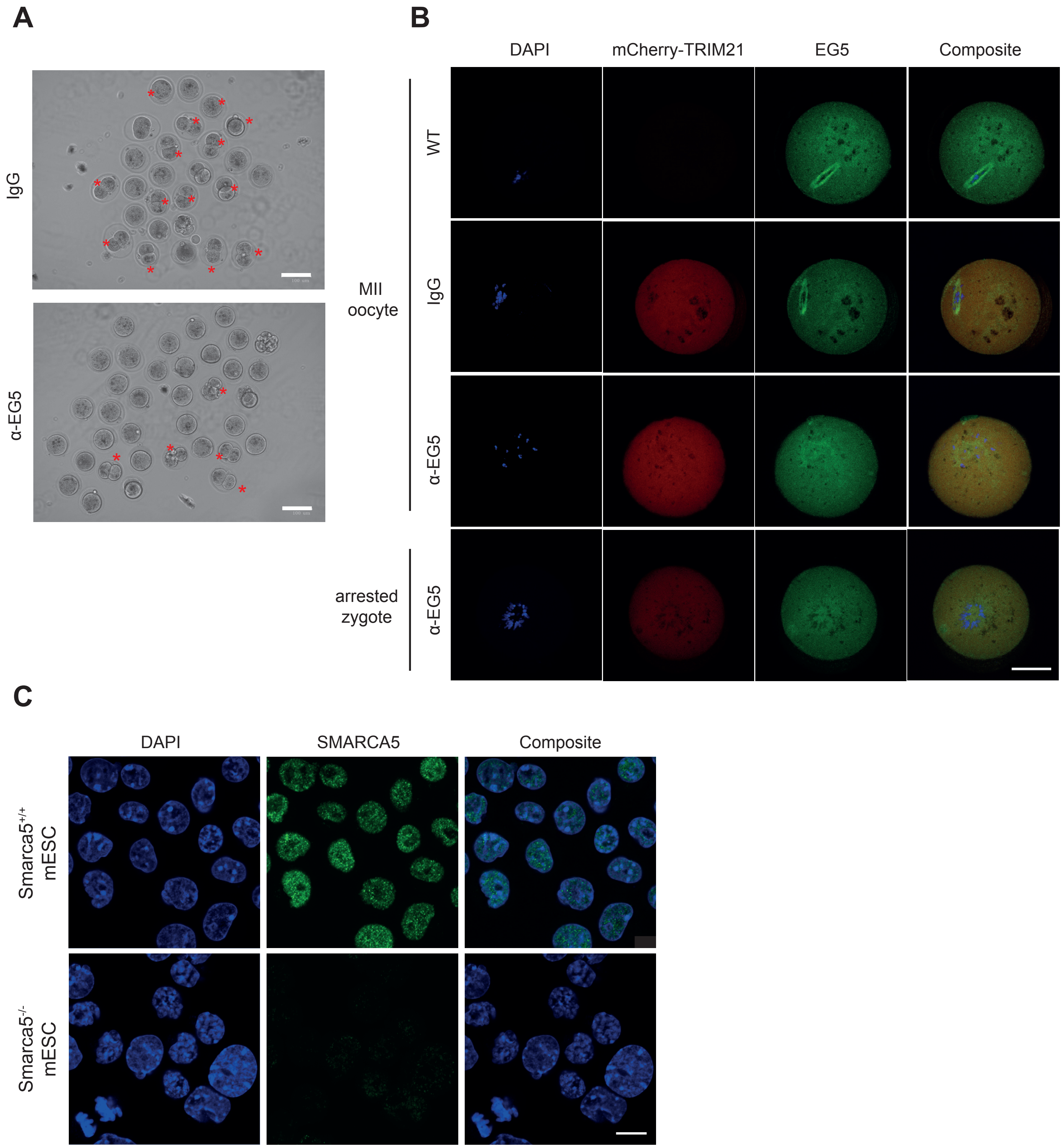


**Supplemental Figure 3**

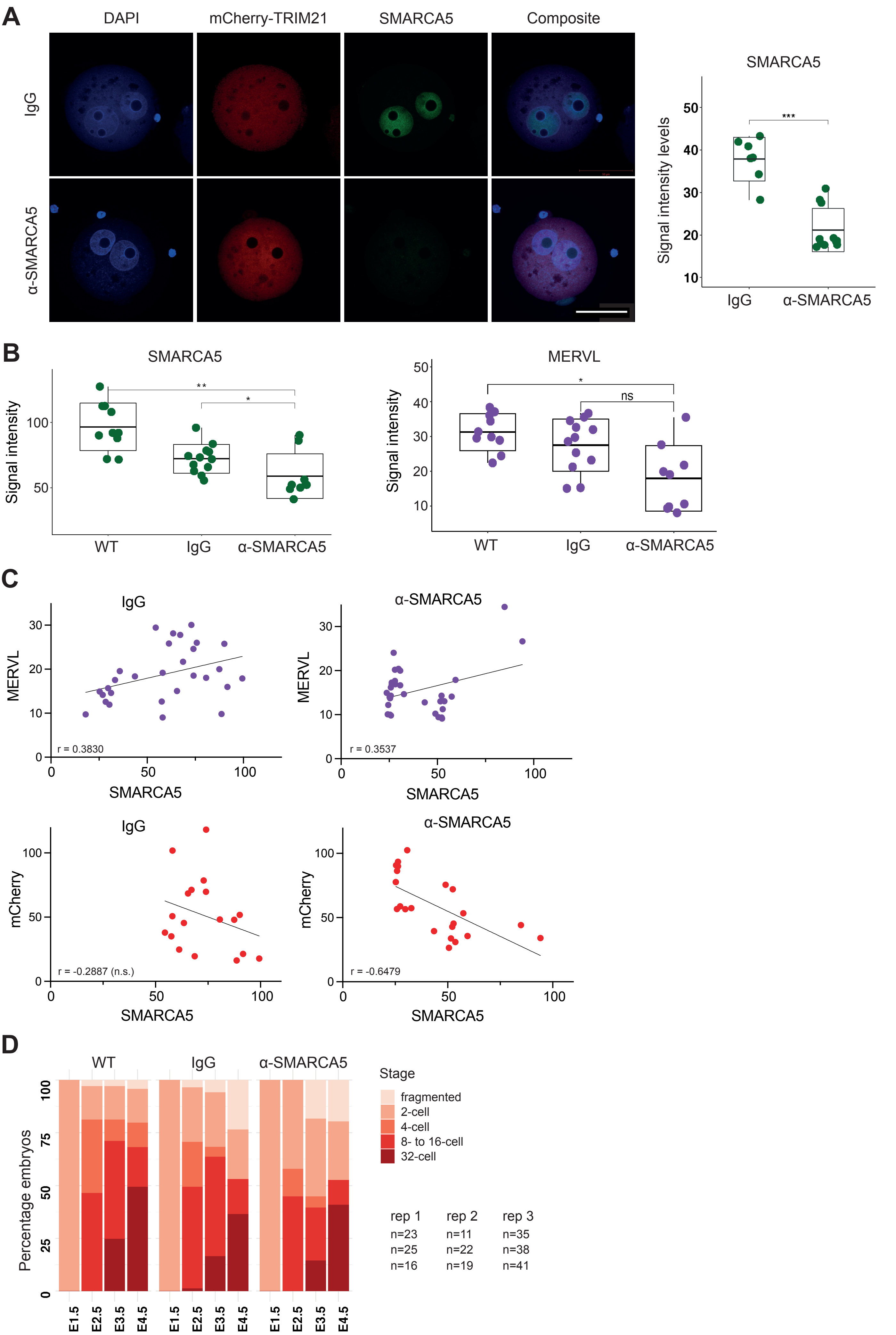


**Supplemental Figure 4**

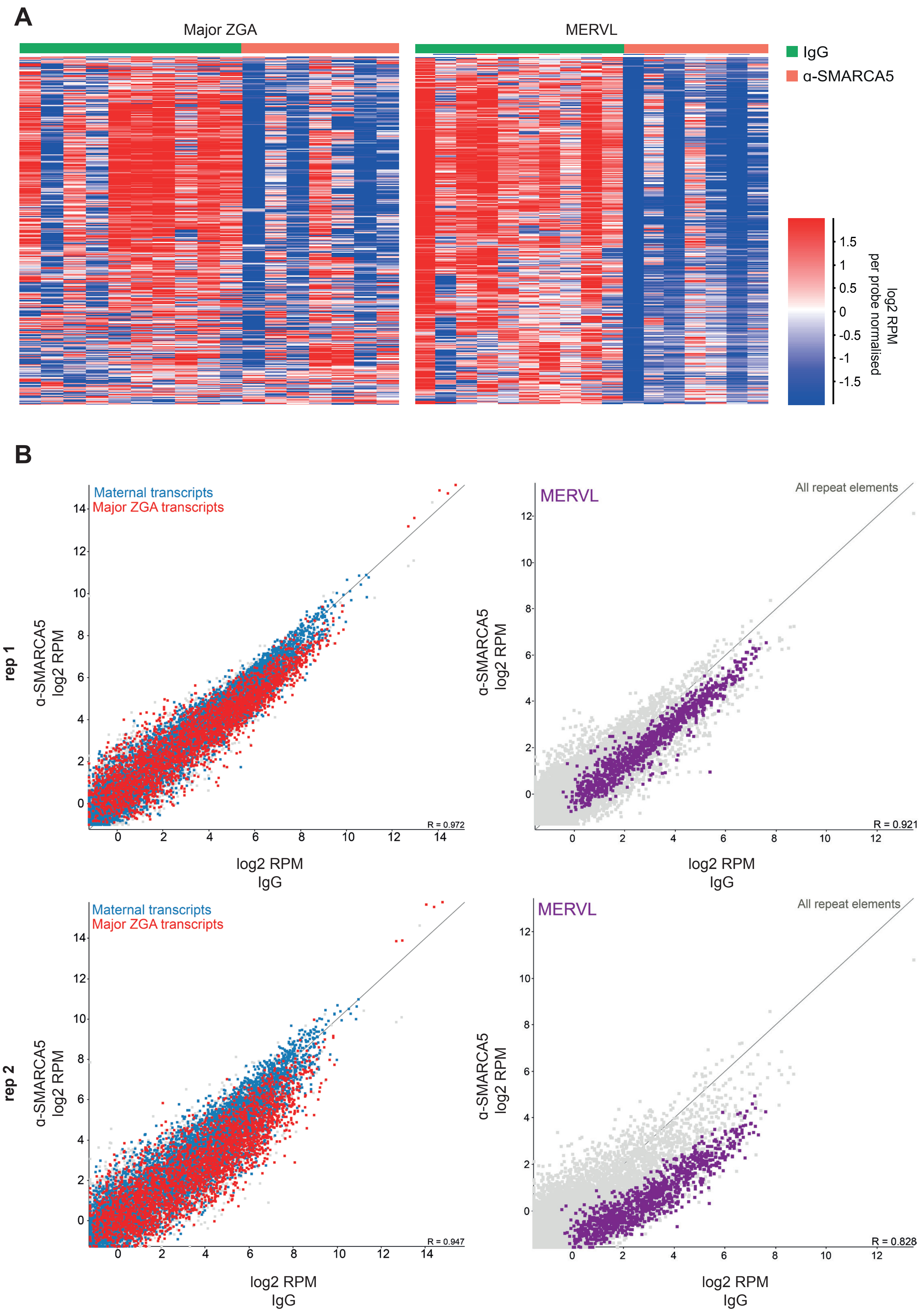


**Supplemental Figure 5**

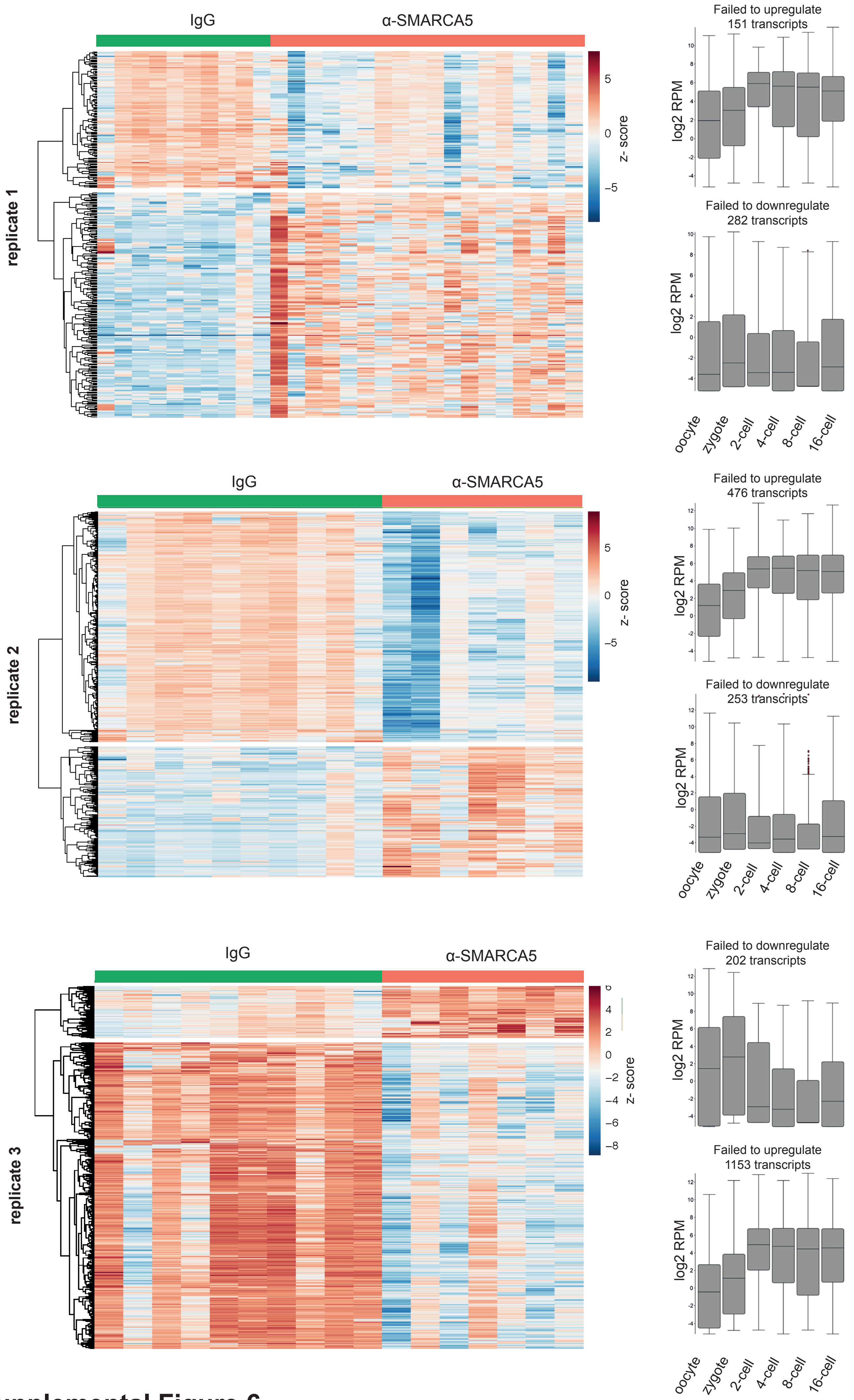


### Supplemental Figure 6

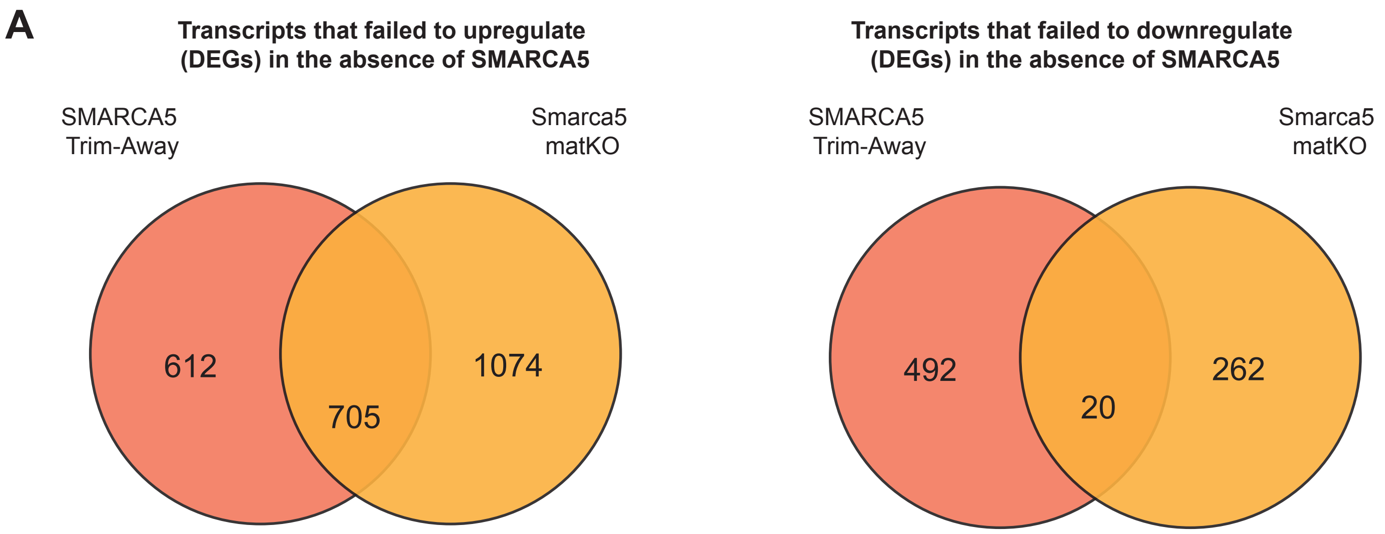


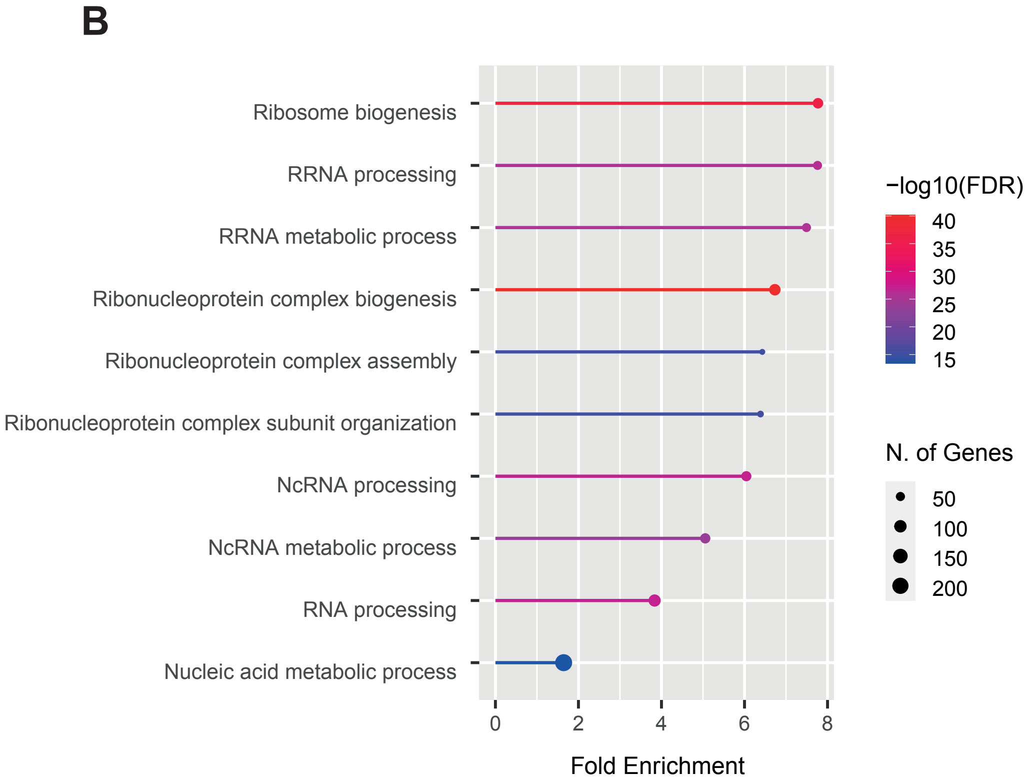


**C**

A630095E13Rik
A730082K24Rik
AU015836
Adra2b
Anxa5
B4galt4
Ceacam20
Fgfr2
Gal
Gm20721
Gm3143
Gm31925
Gm49367
Gnas
Ldhb
Lmo1
Pkd2l2
Prdx2
Tcl1
Tgfb2

#### Supplemental Figure 7

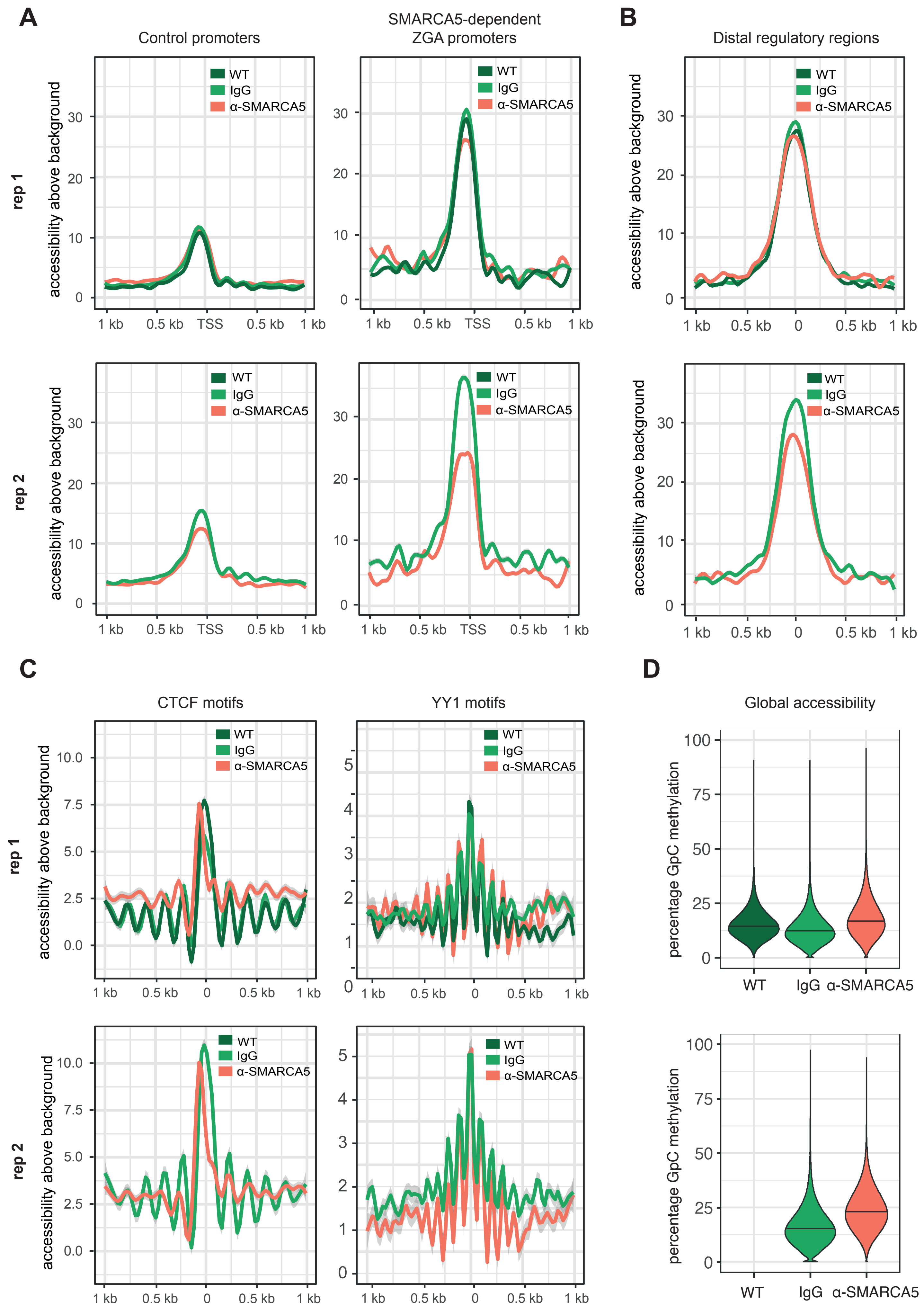


### Supplemental Figure 8

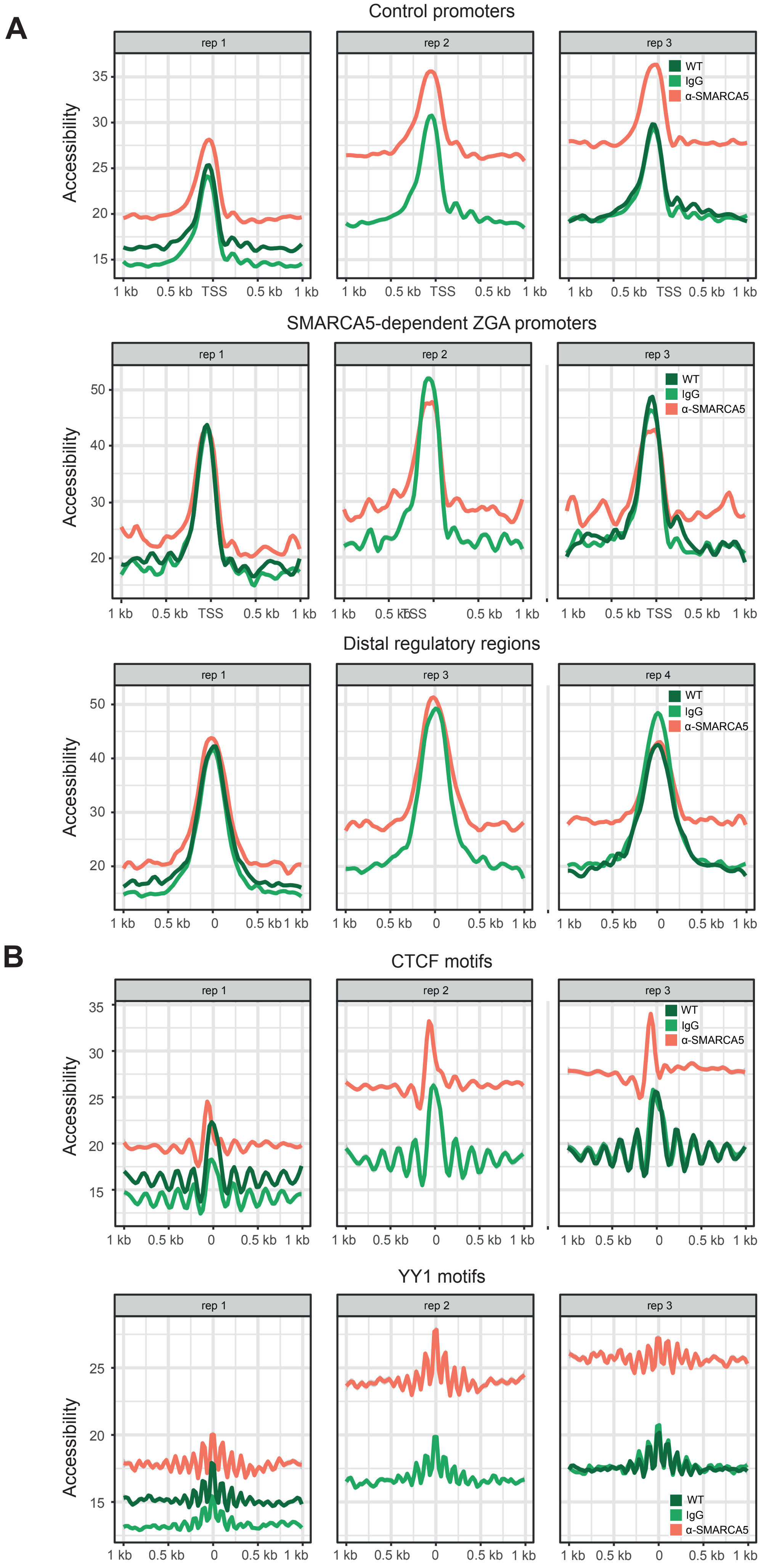


**Supplemental Figure 9**

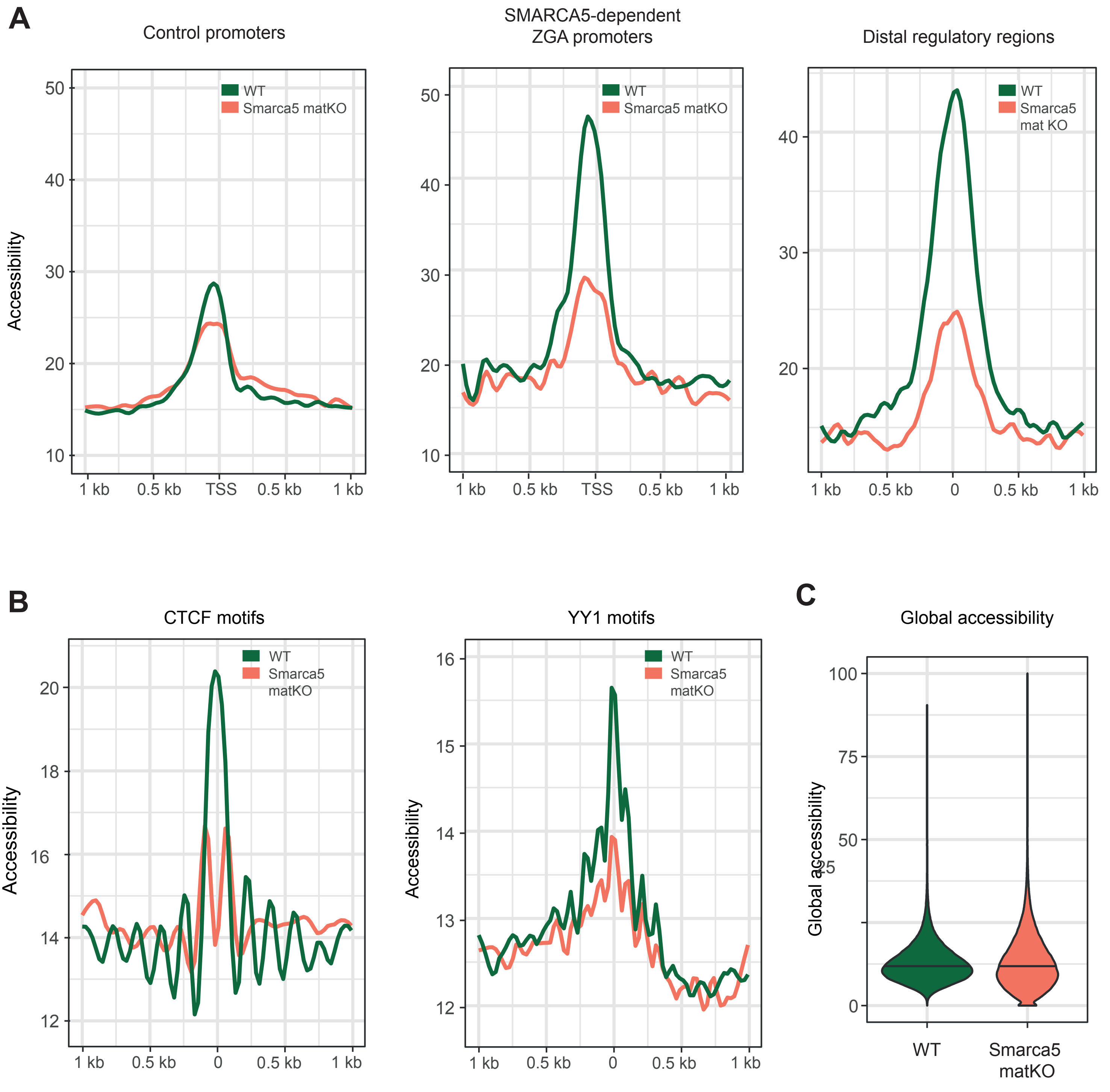


**Supplemental Figure 10**

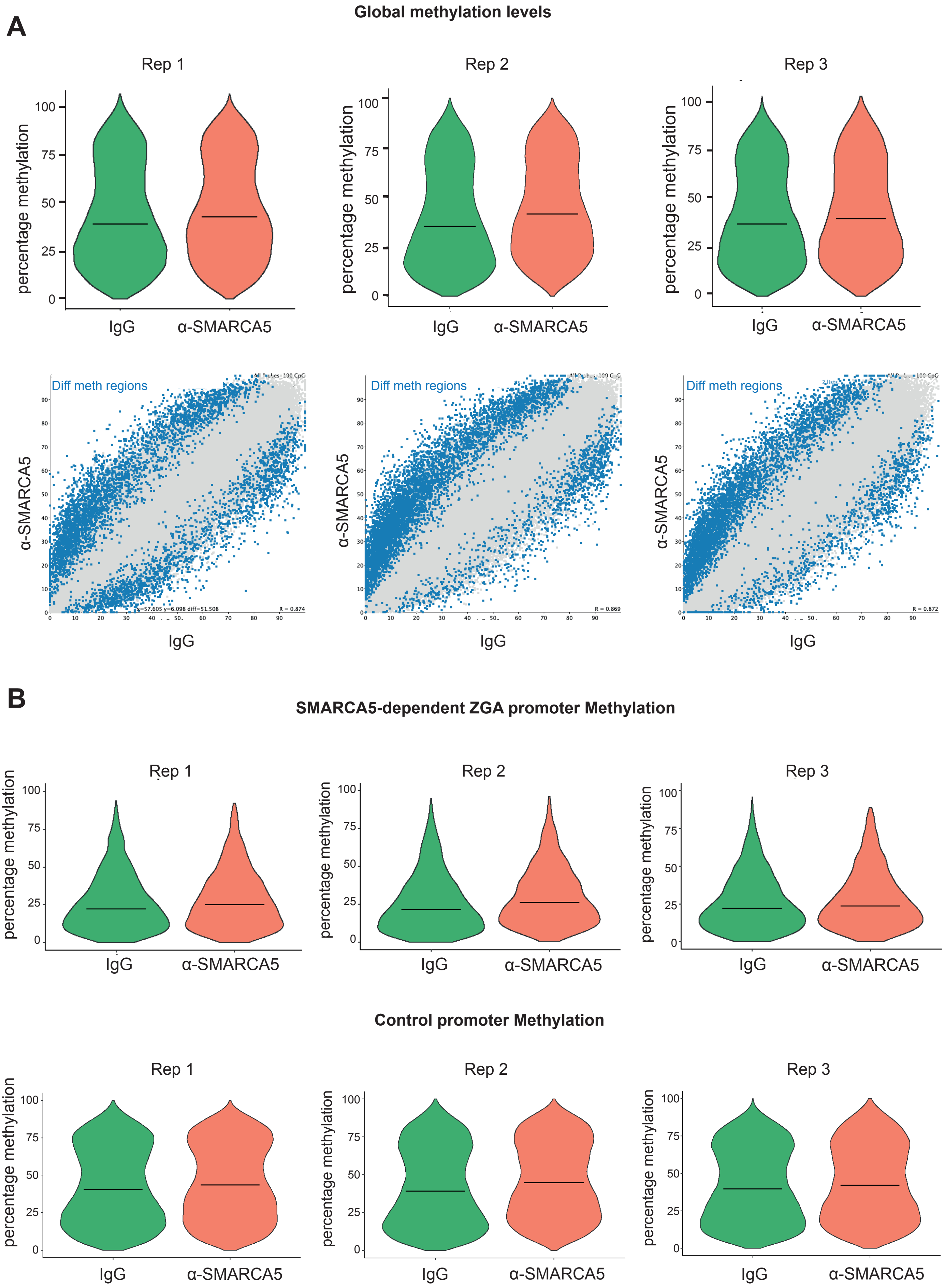


Supplemental Figure 11

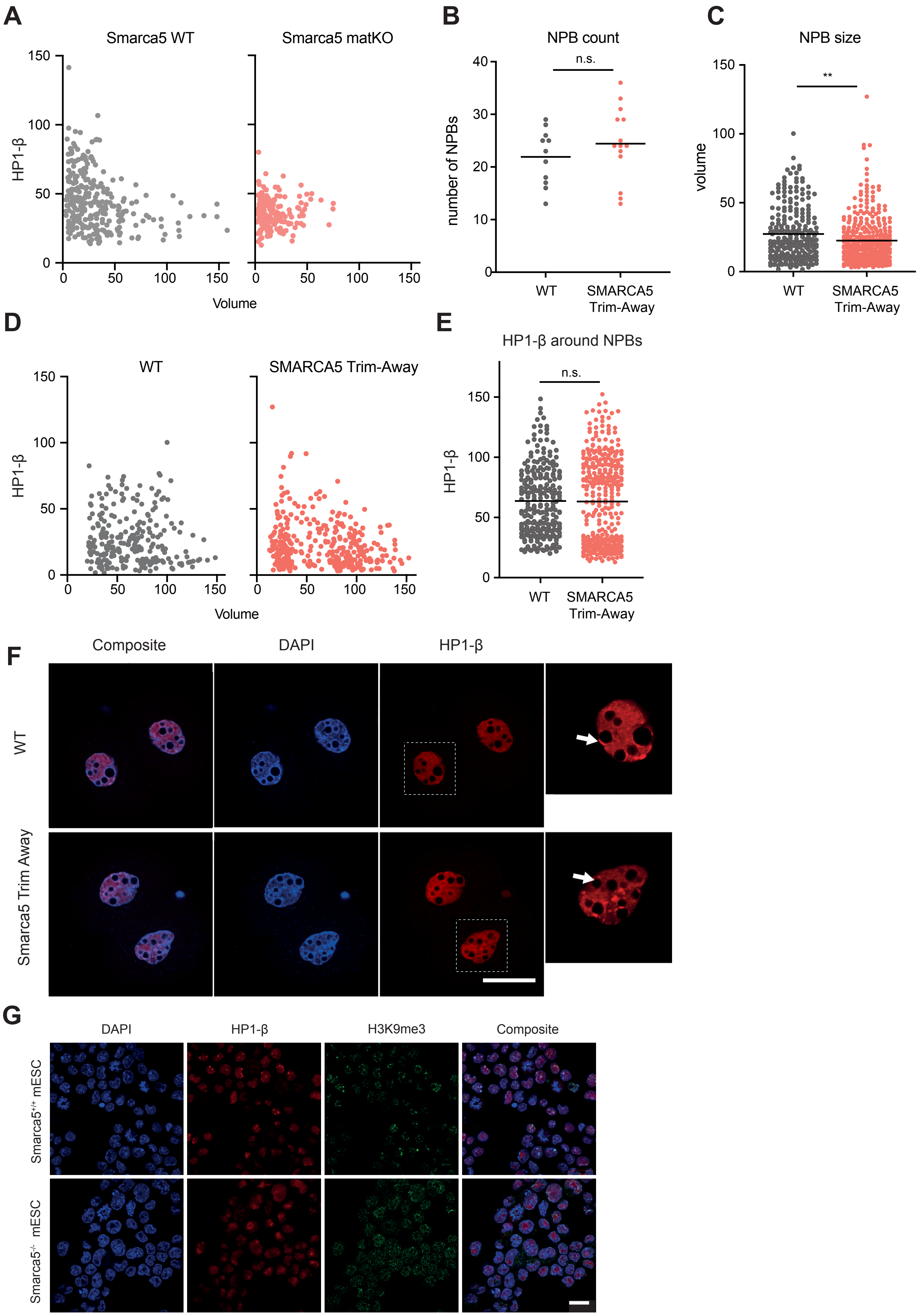


### Supplemental Figure 12
